# Orthogonal neural encoding of targets and distractors supports multivariate cognitive control

**DOI:** 10.1101/2022.12.01.518771

**Authors:** Harrison Ritz, Amitai Shenhav

## Abstract

The complex challenges of our mental life require us to coordinate multiple forms of neural information processing. Recent behavioral studies have found that people can coordinate multiple forms of attention, but the underlying neural control process remains obscure. We hypothesized that the brain implements multivariate control by independently monitoring feature-specific difficulty and independently prioritizing feature-specific processing. During fMRI, participants performed a parametric conflict task that separately tags target and distractor processing. Consistent with feature-specific monitoring, univariate analyses revealed spatially segregated encoding of target and distractor difficulty in dorsal anterior cingulate cortex. Consistent with feature-specific attentional priority, a novel multivariate analysis (Encoding Geometry Analysis) revealed overlapping, but orthogonal, representations of target and distractor coherence in intraparietal sulcus. Coherence representations were mediated by control demands and aligned with both performance and frontoparietal activity, consistent with top-down attention. Together, these findings provide evidence for the neural geometry necessary to coordinate multivariate cognitive control.

## Introduction

We have remarkable flexibility in how we think and act. This flexibility is enabled by the array of mental tools we can bring to bear on challenges to our goal pursuit (Badre et al., 2021; Danielmeier et al., 2011; Egner, 2008; Friedman and Miyake, 2017; Musslick et al., 2015; Ritz et al., 2022a). For example, someone may respond to a mistake by becoming more cautious, enhancing task-relevant processing, or suppressing task-irrelevant processing (Danielmeier and Ullsperger, 2011), and previous work has shown that people simultaneously deploy multiple such strategies at the same time in response to different task demands (Danielmeier et al., 2011; Fischer et al., 2018; Leng et al., 2021; Ritz and Shenhav, 2021). Flexibly coordinating multiple cognitive processes requires a control system that can monitor multiple forms of task demands and deploy multiple forms of control (also referred to as the necessity for *observability* and *controllability*; (Kalman, 1960)). These monitoring and regulation processes are fundamental to control, and are thought to be underpinned by distinct cingulo-opercular and frontoparietal neural systems (Gordon et al., 2017; Gottlieb et al., 2020; Gratton et al., 2016; Kerns et al., 2004; MacDonald et al., 2000; Menon and D’Esposito, 2021; Shenhav et al., 2013; Smith et al., 2019). However, much is still unknown about how multiple forms of control are represented across these domains.

Past research on the neural mechanisms of cognitive control has often sought to identify representations that integrate over multiple different sources of task demands (i.e., represent these different sources in *alignment*). For instance, previous studies has proposed that dorsal anterior cingulate cortex (dACC) tracks integrative features like response conflict, effort, value, error likelihood, and time-on-task (Brown and Braver, 2005; Fu et al., 2022; Grinband et al., 2011; Kragel et al., 2018; Mumford et al., 2023; Rushworth and Behrens, 2008; Vermeylen et al., 2020; Yarkoni et al., 2009). Because they integrate over different task features instead of differentiating between them, these forms of ‘aligned encoding’ (Figure 1a) are ill-suited for carrying out multidimensional control. Multidimensional cognitive control requires independent representations that can track multiple sources of difficulty and regulate multiple cognitive processes (e.g., prioritize multiple sources of information (Pylyshyn and Storm, 1988)).

**Figure 1.**
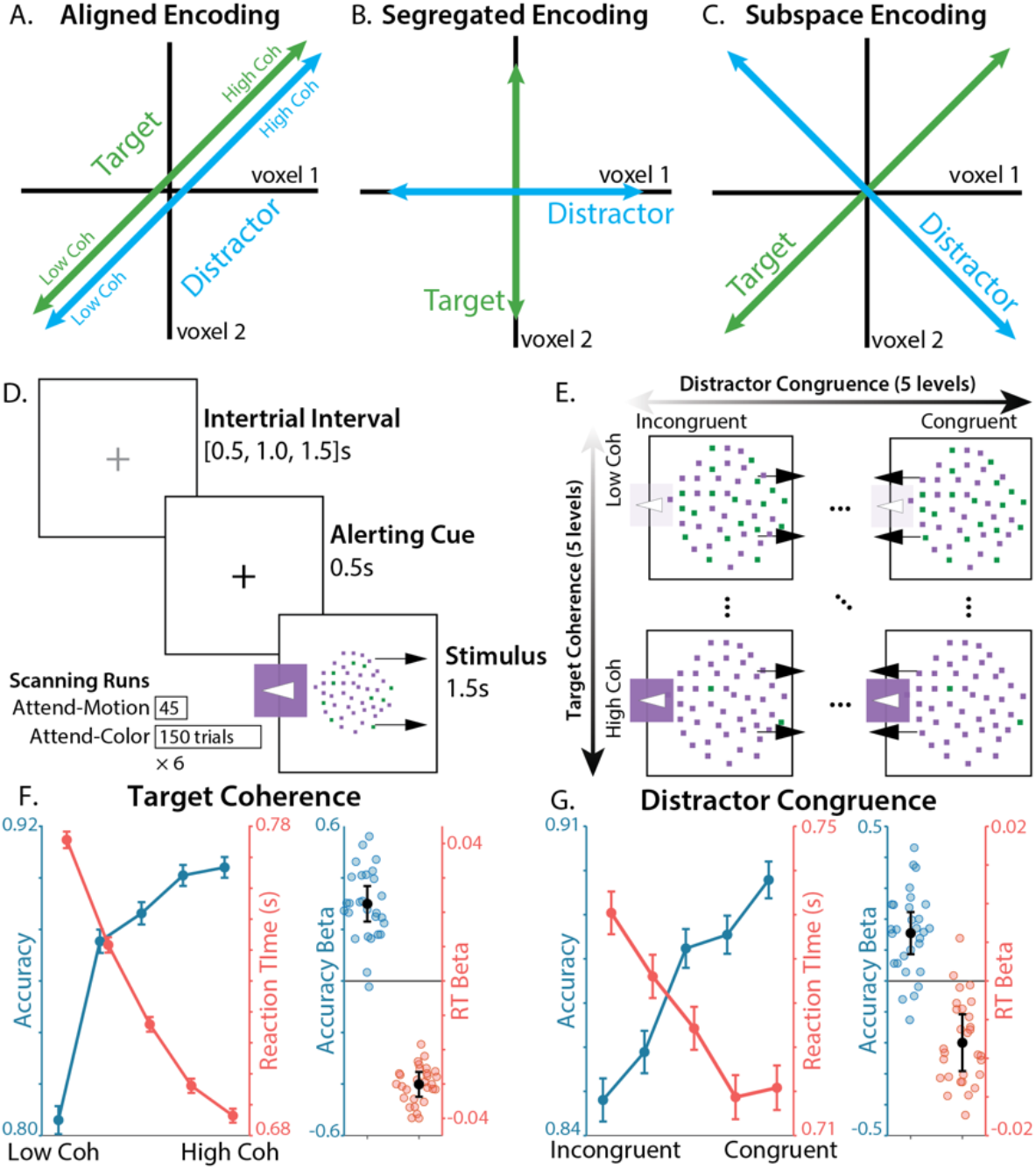
*Task and Behavior.* **A-C)** Three hypothesized encoding schemes. **A)** In *aligned encoding* features are represented similarly, e.g., encode performance variables like error likelihood or time-on-task. **B)** In *segregated encoding* features are encoded independently, in distinct voxel populations (i.e., voxel-level pure selectivity (Rigotti et al., 2013)). **C)** In *subspace encoding,* features are encoding independently, in overlapping voxel populations (i.e., voxel-level mixed selectivity). **D)** Participants responded to a color-motion random dot kinematogram (RDK) with a button press. Participants either responded to the left/right motion direction of the RDK (Attend-Motion runs) or based on the majority color (Attend-Color runs; critical condition). **E)** We parametrically and independently manipulated target coherence (% of dots in the majority color) and distractor congruence (motion coherence signed relative to the target response). **F)** Participants were faster and more accurate when the target was more coherent. **G)** Participants were faster and more accurate when the distractor was more congruent with the target. Error bars on line plots reflect within-participant SEM, error bars for regression fixed-effect betas reflect 95% CI.

An alternative to aligned encoding – one that would allow the brain to separately control multiple processes – is *independent* encoding, which can come in at least two forms. One way the brain can have independent representations is by encoding different task features in spatially segregated neural populations (‘segregated encoding’; Figure 1b). For example, past work has shown that different subregions within dACC encode distinct task demands, including various forms of errors and processing conflict (Beldzik and Ullsperger, 2023; Fu et al., 2019; Shenhav et al., 2018; Taren et al., 2011; Venkatraman et al., 2009; Zarr and Brown, 2016). The brain can instead have independent representations that are distributed across units within the same population, as has also been observed in dACC (Ebitz et al., 2020; Flesch et al., 2022; Minxha et al., 2020). Within a shared population, independent encoding of information occurs along a set of orthogonal dimensions or *subspaces* (Figure 1c, ‘subspace encoding’; (Cunningham and Yu, 2014; Ebitz and Hayden, 2021; Mante et al., 2013; Rigotti et al., 2013)). Despite this exciting recent work, it remains unclear to what extent different components of the cognitive control system leverage these aligned, segregated, or orthogonal encoding strategies for monitoring multiple task demands and prioritizing multiple sources of information.

To gain new insight into the representations supporting cognitive control, we drew upon two key innovations. First, we leveraged an experimental paradigm we developed to tag multiple control processes (Ritz and Shenhav, 2021). Building on prior work (Danielmeier et al., 2011; Kayser et al., 2010b; Mante et al., 2013; Shenhav et al., 2018), this task incorporates elements of perceptual decision-making (discrimination of a target feature) and inhibitory control (overcoming a salient and prepotent distractor). We have previously shown that we can separately tag target and distractor processing from participants’ performance on this task, and that target and distractor processing are independently controlled. For example, participants adjust target and distractor sensitivity in response to distinct task demands (e.g., previous conflict or incentives; (Ritz and Shenhav, 2021)). In conjunction with this process-tagging approach, our second innovation was to develop a novel multivariate fMRI analysis for measuring relationships between feature encoding (i.e., *encoding geometry*). Extending recent statistical approaches in systems neuroscience (Bernardi et al., 2020; Ebitz et al., 2020; Panichello and Buschman, 2021), we combined the strengths of multivariate encoding analyses and representation similarity analyses into a method we call ‘Encoding Geometry Analysis’ (EGA). We used EGA to characterize whether putative markers of monitoring and prioritization leverage independent representations for targets and distractors.

In brief, we found that key nodes within the cognitive control network use orthogonal representations of target and distractor information to support cognitive control. In the dorsal anterior cingulate cortex (dACC), encoding of target and distractor difficulty was spatially segregated and arranged along a rostrocaudal gradient. By contrast, in the intraparietal sulcus (IPS), encoding of target and distractor coherence was encoded along orthogonal neural subspaces. These regional distinctions are consistent with hypothesized roles in planning and implementing (multivariate) attentional policies (Gottlieb et al., 2020; Shenhav et al., 2013). Furthermore, we found that coherence encoding depended on control demands, and was aligned with both task performance and frontoparietal activity, consistent with these coherence representations playing a critical role in cognitive control (e.g., feature prioritization). Together, these results suggest that cognitive control uses representational formats that allow the brain to monitor and control multiple streams of information processing.

## Results

### Task overview

Twenty-nine human participants performed the Parametric Attentional Control Task (PACT; (Ritz and Shenhav, 2021) during fMRI. On each trial, participants responded to an array of colored moving dots (colored random dot kinematogram; Figure 1d). In the critical condition (Attend-Color), participants respond with a left/right keypress based on which of two colors were in the majority. In alternating scanner runs, participants instead responded based on motion (Attend-Motion), which was designed to be less control-demanding due to the (Simon-like) congruence between motion direction and response hand (Danielmeier et al., 2011; Ritz and Shenhav, 2021). Across trials, we independently and parametrically manipulated target and distractor information across five levels of target coherence (e.g., percentage of dots in the majority color, regardless of which color) and distractor congruence (e.g., percentage of dots moving either in the congruent or incongruent direction relative to the correct color response; Figure 1e). This task allowed us to ‘tag’ participants’ sensitivity to each dimension by measuring behavioral and neural responses to independently manipulated target and distractor features. Unlike a similar task used to study post-error adjustments (Danielmeier et al., 2011), our parametric manipulation of target and distractor coherence allows us to better measure feature-specific representations. Unlike similar tasks used to study contextual decision-making (Mante et al., 2013; Shenhav et al., 2018; Takagi et al., 2021), this task pits more control-demanding responses (towards color) against more automatic responses (towards motion), allowing comparisons between Attend-Color and Attend-Motion tasks to isolate the contributions of cognitive control (Cohen et al., 1990; Woolgar et al., 2011a).

### Behavior

Participants had overall good performance on the task, with a high level of accuracy (median Accuracy = 89%, IQR = [84% - 92%]), and a low rate of missed responses (median lapse rate = 2%, IQR = [0% - 5%]). We used mixed effects regressions to characterize how target coherence and distractor congruence influenced participants’ accuracy and log-transformed correct reaction times. Replicating previous behavioral findings using this task, participants were sensitive to both target and distractor information (Ritz and Shenhav, 2021). When target coherence was weaker, participants responded slower (*t*(27.6) = 16.1, *p* = 1.60 × 10^-15^) and less accurately (*t*(28) = - 8.90, *p* = 1.19 × 10^-9^; Figure 1f). When distractors were more incongruent, participants also responded slower (*t*(28.8) = 5.09, *p* = 2.15 × 10^-5^) and less accurately (*t*(28) = -4.66, *p* = 6.99 × 10^-5^; Figure 1g). Also replicating prior findings with this task, interactions between targets and distractors were not significant for reaction time (*t*(28.2) = 0.143, *p* = .887) and had a weak influence on accuracy (*t*(28) = 2.36, *p* = .0257), with model omitting target-distractor interactions providing a better complexity-penalized fit (RT ΔAIC = 17.7, Accuracy ΔAIC = 1.38).

### Segregated encoding of target and distractor difficulty in dACC

Past work has separately shown that the dACC tracks task demands related to perceptual discrimination (induced in our task when target information is weaker) and related to the need to suppress a salient distractor (induced in our task when distractor information is more strongly incongruent with the target (Nee et al., 2007; Shenhav et al., 2018, 2013; Taren et al., 2011; Venkatraman et al., 2009)). Our task allowed us to test whether these two sources of increasing control demand are tracked within common regions of dACC (reflecting an aggregated representation of multiple sources of task demands), or whether they are tracked by separate regions (potentially reflecting a specialized representation according to the nature of the demands).

Targeting a large region of dACC – a conjunction of a cortical parcellation with a meta-analytic mask for ‘cognitive control’ (see ‘fMRI univariate analyses’ in Methods) – we found spatially distinct signatures of target difficulty and distractor congruence within dACC. In caudal dACC, we found significant clusters encoding the parametric effect of target difficulty (Figure 2a; negative effect of target coherence in green), and in more rostral dACC we found clusters encoding parametric distractor incongruence (negative effect of distractor congruence in blue). Supporting this dissociation, the spatial patterns of target and distractor regression weights were uncorrelated across dACC voxels (*t*(28.0) = 1.32, *p* = .197, logBF = -0.363). These analyses control for omission errors, and additionally controlling for commission errors produced the same whole-brain pattern at a reduced threshold (see Supplementary Figure 1). While distractor congruence was marginally significant when correcting for multiple comparisons across the entire brain (one-sided *p* = .08, whole-brain TFCE), extensive previous research predicts congruence effects in this ROI and in this direction (Kragel et al., 2018; Nee et al., 2007; Shenhav et al., 2013), suggesting that a whole-brain corrected estimate is overly conservative.

**Figure 2.**
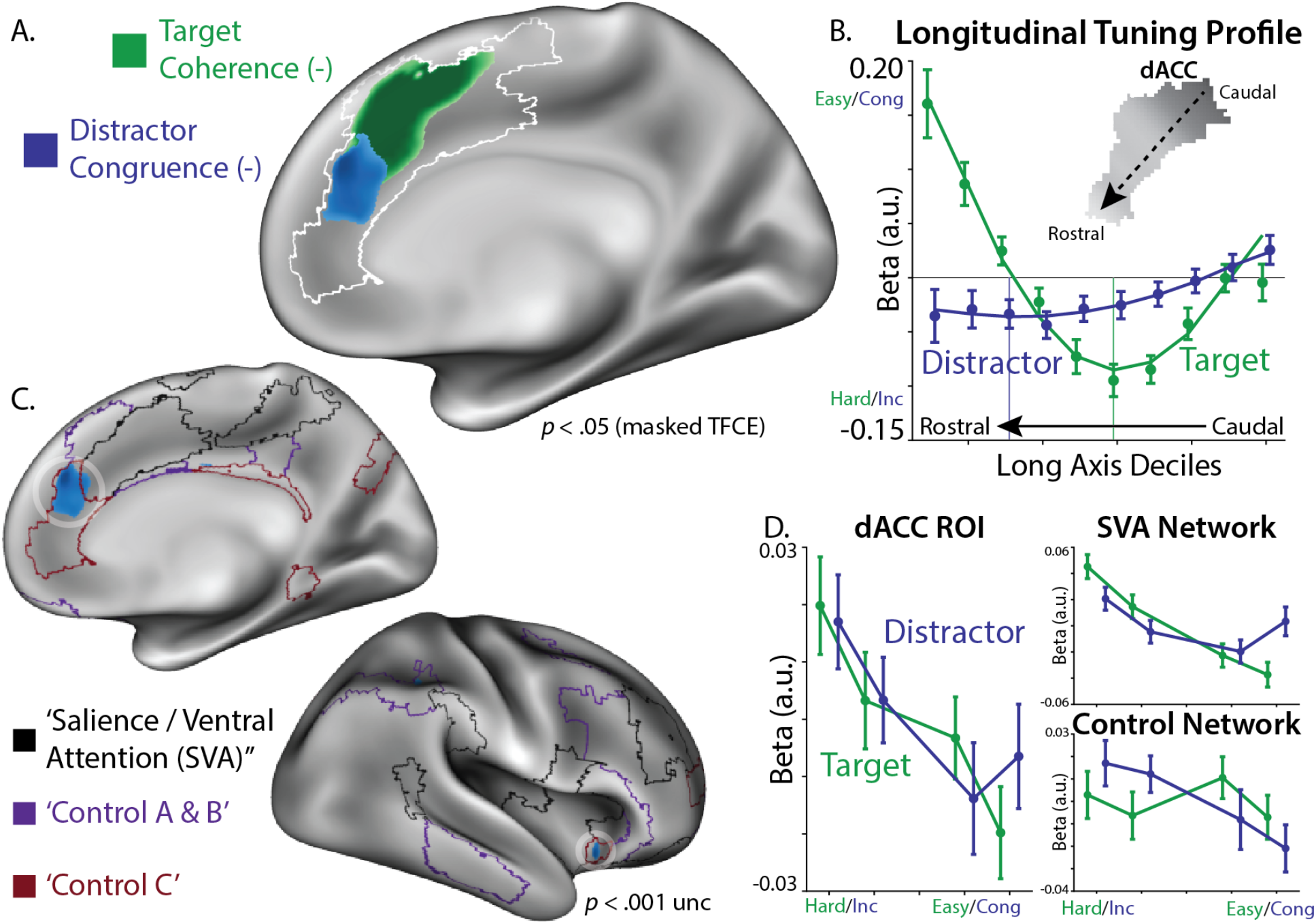
*Distinct coding of target and distractor difficulty in dACC.* **A)** We looked for linear target coherence and distractor congruence signals within an a priori dACC mask (white outline; overlapping Kong22 parcels and medial ‘cognitive control’ Neurosynth mask). We found that voxels in the most caudal dACC reflected target difficulty (green), more rostral voxels reflected distractor incongruence (blue). Statistical tests are corrected using non-parametric threshold-free cluster enhancement. **B)** We extracted the long axis of the dACC using a PCA of the voxel coordinates. We plotted the target coherence (green) and distractor congruence (blue) along the deciles of this long axis. Fit lines are the quantized predictions from a second-order polynomial regression. We used these regression betas to estimate the minima for target and distractor tuning (i.e., location of strongest difficulty effects), finding that the target difficulty peak (vertical green line) was more caudal than the distractor incongruence peak (vertical blue line). **C)** Plotting the uncorrected whole-brain response, distractor incongruence responses (blue) were strongest within the ‘Control C’ sub-network (red), both in dACC and anterior insula. **D)** BOLD responses across levels of target coherence and distractor congruence, plotted within the whole dACC ROI (left), or the ‘Salience/Ventral Attention (SVA)’ network and ‘Control’ network parcels within the dACC ROI (right). GLMs: A-C: Feature UV, D: Difficulty Levels, see Table 2.

**Table 2.** *fMRI models*. First-level general linear models used for univariate and multivariate fMRI analyses. Coherence: percentage of dots supporting the same response (’unsigned coherence’). Evidence: % dots supporting a rightwards vs leftwards response (‘signed coherence’). Distractor Congruence: % dots supporting the same response as the target dimension. All predictors were z-scored within their run. For difficulty and feature levels, we included each level as a separate predictor, with collinearity with the block intercept preventing all levels from being included. For Evidence Levels, targets had greater granularity due to distractors being coded relative to targets (5 levels of congruence led to 5 levels of coherence). For Performance CX, seed timeseries were included as run-separated regressors (see Multivariate Connectivity Analysis in Methods).

| Model Name | Trial selection | Predictors (z-scored) |
| --- | --- | --- |
| Feature UV | No omission errors;<br>run-concatenated | target coherence, distractor coherence, target evidence, distractor evidence, distractor congruence;<br>omission errors (run-concatenated) |
| Difficulty Levels | No omission errors;<br>run-concatenated | Separate levels (1,2,4,5) of target coherence, separate levels (1,2,4,5) of distractor congruence;<br>omission errors (run-concatenated) |
| Feature MV | No errors;<br>run-separated | target coherence, distractor coherence, target evidence, distractor evidence, distractor congruence;<br>errors (run-concatenated) |
| Evidence Levels | No errors;<br>run-separated | Levels (1-5, 6-10) of target evidence, Levels (1,2,4,5) of distractor evidence;<br>errors (run-concatenated) |
| Between-Task | No errors;<br>run-separated | target coherence, distractor coherence, target evidence, distractor evidence, distractor congruence;<br>errors (run-concatenated);<br>reaction time (run-concatenated) |
| Performance | No omission errors;<br>run-separated | target coherence, distractor coherence, target evidence, distractor evidence, distractor congruence, reaction time, accuracy;<br>omission errors (run-concatenated) |
| Performance CX | No omission errors;<br>run-separated | target coherence, distractor coherence, target evidence, distractor evidence, distractor congruence, reaction time, accuracy;<br>omission errors (run-concatenated);<br>seed timeseries (run-separated) |

To further quantify how feature encoding changed along the longitudinal axis of dACC, we used principal component analysis to extract the axis position of dACC voxels (see ‘dACC longitudinal axis analyses’ in Methods), and then regressed target and distractor beta weights onto these axis scores. We found that targets had stronger difficulty coding in more caudal voxels (*t*(27.9) = 3.74, *p* = .000840), with a quadratic trend (*t*(26.5) = 4.48, *p* = .000129; Figure 2b). In line with previous work on both perceptual and value-based decision-making (Clairis and Pessiglione, 2022; Fleming et al., 2018; Shenhav et al., 2018, 2016; Shenhav and Karmarkar, 2019), we found that signatures of target discrimination difficulty (negative correlation with target coherence) in caudal dACC were paralleled by signals of target discrimination *ease* (positive correlation with target coherence) within the rostral-most extent of our dACC ROI (Supplementary Figure 2). In contrast to targets, distractors had stronger incongruence coding in more rostral voxel (*t*(28.0) = -3.26, *p* = .00294), without a significant quadratic trend. We used participants’ random effects terms to estimate the gradient location where target and distractor coding were at their most negative, finding that the target minimum was significantly more caudal than the distractor minimum (signed-rank test, *z*(28) = 2.41, *p* = .0159). Target and distractor minima were uncorrelated across subjects (*r*(27) = .0282, *p* = .880, logBF = -0.839), again consistent with independent encoding of targets and distractors.

As additional evidence that target-related and distractor-related demands have a dissociable encoding profile, we found that the crossover between target and distractor encoding in dACC occurred at the boundary between two well-characterized functional networks (Kong et al., 2021; Schaefer et al., 2018; Yeo et al., 2011). Whereas distractor-related demands were more strongly encoded rostrally in the Control Network (particularly within regions of dACC and insula corresponding to the ‘Control C’ Sub-Network; (Kong et al., 2021, 2019)), target-related demands were more strongly encoded caudally within the ‘Salience / Ventral Attention (SVA)’ Network (Figure 2C-D). Including network membership alongside long axis location predicted target and distractor encoding better than models with either network membership or axis location alone (*Δ*BIC > 1675).

### Subspace encoding of target and distractor coherence in intraparietal sulcus

We found that dACC appeared to dissociably encode target and distractor difficulty through spatially segregated encoding, consistent with a role in monitoring different task demands and/or specifying different control signals (Shenhav et al., 2013). To identify neural mechanisms for the implementation of this control through the prioritization targets versus distractors, we next tested for regions that encode target and distractor coherence (the amount of information in a feature, regardless of which response it supports). Based on previous research, we might expect to find this form of selective attention in posterior parietal cortex (Bisley and Mirpour, 2019; Gottlieb et al., 2020; Yantis and Serences, 2003). We explored whether target and distractor coherence share a common neural code (e.g., as a global index of spatial salience), compared to where these features are encoded distinctly (e.g., as separate targets of control).

An initial whole-brain univariate analysis showed that overlapping regions throughout occipital, parietal, and prefrontal cortices track the feature coherence (proportion of dots in the majority category) for both targets and distractors (Figure 3a; conjunction in orange). These regions showed elevated responses to lower target coherence and higher distractor coherence, potentially reflecting the relevance of each feature for task performance. Note that in contrast to distractor congruence, distractor *coherence* had an inconsistent relationship with task performance (RT: *t*(27.0) = 2.08, *p* = .048; Accuracy: *t*(28) = -0.845, *p* = .406), suggesting that these neural responses are unlikely to reflect task difficulty per se.

**Figure 3.**
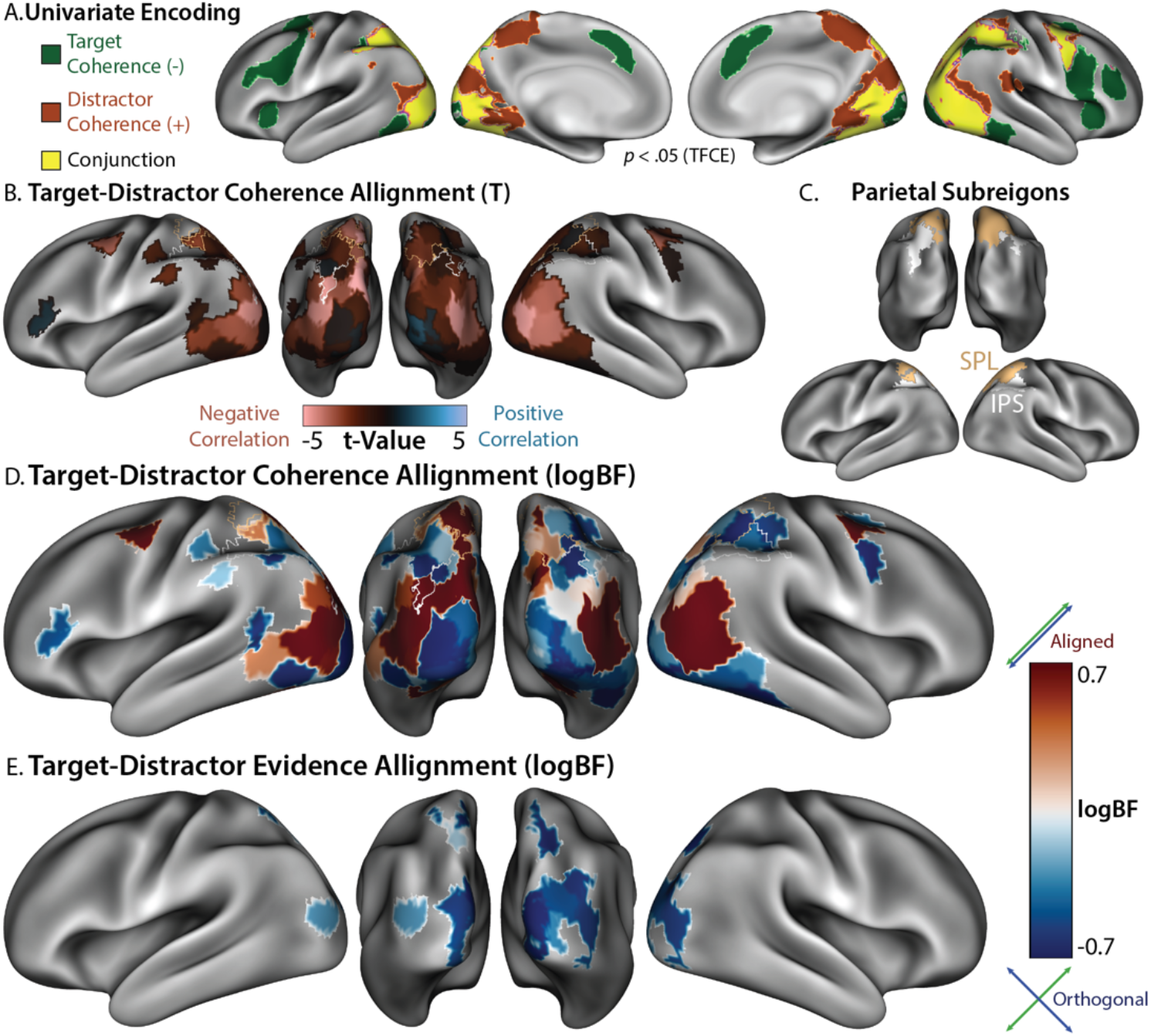
*Encoding Geometry Analysis (EGA) dissociates target and distractor encoding.* **A)** Parametric univariate responses to weak target coherence (green; percentage of dots in majority color), strong distractor coherence (orange; percentage of dots with coherent motion), and their conjunction (yellow). Statistical tests are corrected for multiple comparisons using non-parametric threshold-free cluster enhancement (TFCE). **B)** Encoding alignment within parcels in which target and distractor encoding was jointly reliable (both *p* < .001 uncorrected). Representations were negatively correlated within Superior Parietal Lobule (SPL in gold; Kong22 labels), and uncorrelated within Intraparietal Sulcus (IPS in white; Kong22 labels). **C)** Anatomical labels for parietal regions, based on the labels in the Kong22 parcellation. **D)** Bayesian analyses provide explicit evidence for orthogonality within IPS (i.e., negative BF; theoretical minima: -0.71). **E)** Coherence coded in terms of evidence (i.e., supporting a left vs right choice). Target and distractor evidence encoding overlapped in visual cortex and SPL and was represented orthogonally. GLMs: A: Feature UV, B-E: Feature MV, see Table 2.

While these univariate activations point towards widespread and coarsely overlapping encoding of the feature coherence (potentially consistent with aligned encoding; Figure 1a), they lack information about how these features are encoded at finer spatial scales. To interrogate the relationship between target and distractor encoding, we developed a multivariate analysis that combines multivariate encoding analyses with pattern similarity analyses, which we term Encoding Geometry Analysis (EGA). Whereas pattern similarity analyses typically quantify relationships between representations of specific stimuli or responses (e.g., whether they could be classified, (Kriegeskorte and Diedrichsen, 2019)), EGA characterizes relationships between encoding subspaces (patterns of contrast weights) across different task features, consistent with recent analyses trends in systems neuroscience (Bernardi et al., 2020; Cohen and Maunsell, 2010; Ebitz et al., 2020; Flesch et al., 2022; Kimmel et al., 2020; Libby and Buschman, 2021). A stronger correlation between encoding subspaces (either positive or negative) indicates that features are similarly encoded (i.e., that their representations are aligned and thus confusable by a decoder; Figure 1a), whereas weak correlation indicate that these representations are orthogonal (and thus distinguishable by a decoder; (Kriegeskorte and Diedrichsen, 2019)). In contrast to standard pattern similarity, the sign of these relationships is interpretable in EGA, reflecting how features are coded relative to one another. Compared to standard encoding analyses, EGA is less sensitive to noise (Supplementary Figure 3). We estimated this encoding alignment within each parcel, correlating unsmoothed and spatially pre-whitened patterns of parametric regression betas across scanner runs to minimize spatiotemporal autocorrelation (Diedrichsen and Kriegeskorte, 2017; Nili et al., 2014; Walther et al., 2016). This cross-validated similarity further allowed us to anchor our analysis on the measurement reliability of encoding profiles (i.e., the self-correlation of encoding patterns across cross-validation folds (Spearman, 1987; Thornton and Mitchell, 2017)).

Focusing on regions that encoded both target and distractor information (parcels where both group-level *p* < .001), EGA revealed clear dissociations between regions that represent these features in alignment versus orthogonally. Within visual cortex and the superior parietal lobule (SPL), target and distractor representations demonstrated significant negative correlations (Figure 3b, red), reflecting (negatively) aligned encoding. In contrast, early visual cortex and intraparietal sulcus (IPS; see Figure 3c for anatomical boundaries) demonstrated target-distractor correlations near zero (Figure 3b, black), suggesting encoding along orthogonal subspaces.

To bolster our interpretation of the latter findings as reflecting orthogonal (i.e., uncorrelated) representations rather than merely small but non-significant correlations, we employed Bayesian t-tests at the group level to estimate the relative (log-10) likelihood that these encoding dimensions were orthogonal or correlated. Consistent with our previous analyses, we found strong evidence for correlation (positive log bayes factors) in more medial regions of occipital and posterior parietal cortex (e.g., SPL), and strong evidence for orthogonality (negative log bayes factors) in more lateral regions of occipital and posterior parietal cortex (e.g., IPS; Figure 3D). Control analyses confirmed that coherence orthogonality was not due to encoding reliability, as a similar topography was observed with disattenuated correlations (normalizing correlations by their reliability; see Supplementary Figure 4). Further supporting these results, our Bayes factor analyses were robust to the choice of priors (see Supplementary Figure 5).

While our analyses support independent encoding of targets and distractors within the same parcel, we further explored whether feature information is reflected in overlapping voxels (i.e., voxel-level mixed selectivity (Rigotti et al., 2013)). Simulations revealed that the alignment between absolute encoding weights can differentiate between pure and mixed selectivity, and parietal coherence representations bore this signature of voxel-level mixed selectivity (Supplementary Figure 6), consistent with the subspace encoding hypothesis.

These results have focused on the coherence of different features regardless of the response they support, demonstrating that SPL exhibits aligned representations of target and distractor coherence. Past decision-making research has separately demonstrated that SPL tracks the amount of evidence supporting specific response (Hunt et al., 2012; Kayser et al., 2010a, 2010b), which we found was also true for our task. In addition to encoding target and distractor coherence, SPL and visual cortex also tracked target and distractor ‘evidence’ (proportion of dots supporting a rightward vs leftward response; Figure 3e). EGA revealed orthogonal evidence representations between targets and distractors, in the same areas with aligned coherence representations (compare Figure 3d and 3e), consistent with previous observations of multiple decision-related signals in SPL (Hunt et al., 2012).

We complemented our whole-brain analyses with ROI analyses in areas exhibiting reliable encoding of key variables, focusing on core frontal regions linked with cognitive control (dACC and lateral PFC [lPFC]), and parietal regions linked with decision-making and attention (SPL and IPS; (Menon and D’Esposito, 2021; Shenhav et al., 2013)). Consistent with our analyses above, we found that target and distractor coherence encoding was aligned in SPL, but not in IPS (Figure 4a, compare to Figure 3d), whereas SPL encoded target and distractor evidence. Directly comparing these regions (see Table 1), we found stronger encoding of target evidence in SPL, stronger encoding of target coherence in IPS, and stronger alignment between target-distractor coherence alignment in SPL. Unlike our univariate results, we did not find distractor congruence encoding in dACC (though this was found in lPFC and IPS). Instead, dACC showed multivariate encoding of target coherence and evidence.

**Figure 4.**
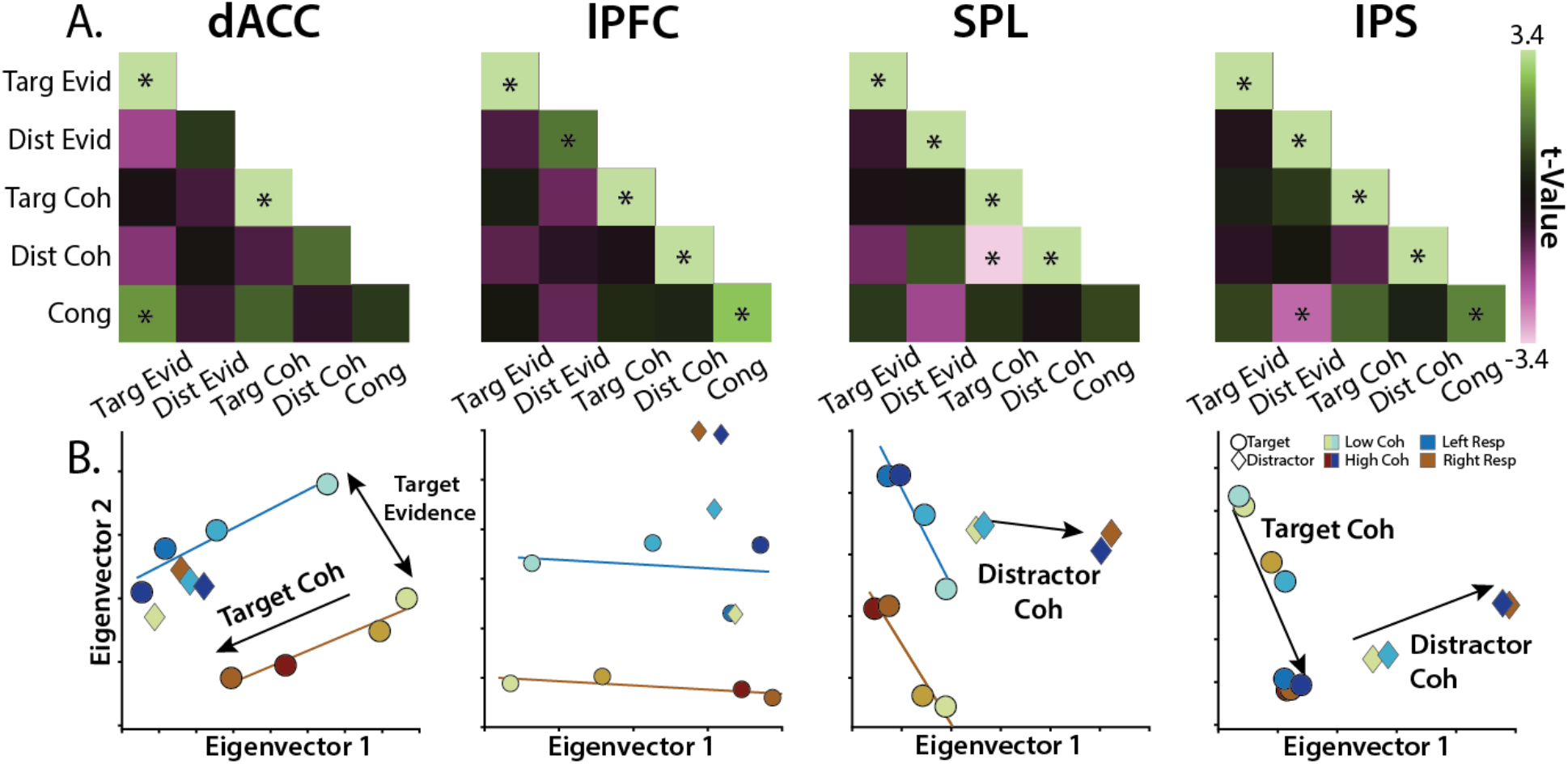
*Region-specific feature encoding.* **A)** Similarity matrices for dACC, lPFC, SPL, and IPS, correlating feature evidence (‘Evid’), feature coherence (‘Coh’), and feature congruence (‘Cong’). Encoding strength on diagonal (right-tailed *p*-value), encoding alignment on off-diagonal (two-tailed *p*-value). **B)** Classical MDS embedding of target (circle) and distractor (diamond) representations at different levels of evidence. Colors denote responses, hues denote coherence. GLMs: A: Feature MV, B: Evidence Levels, see Table 2.

**Table 1.**
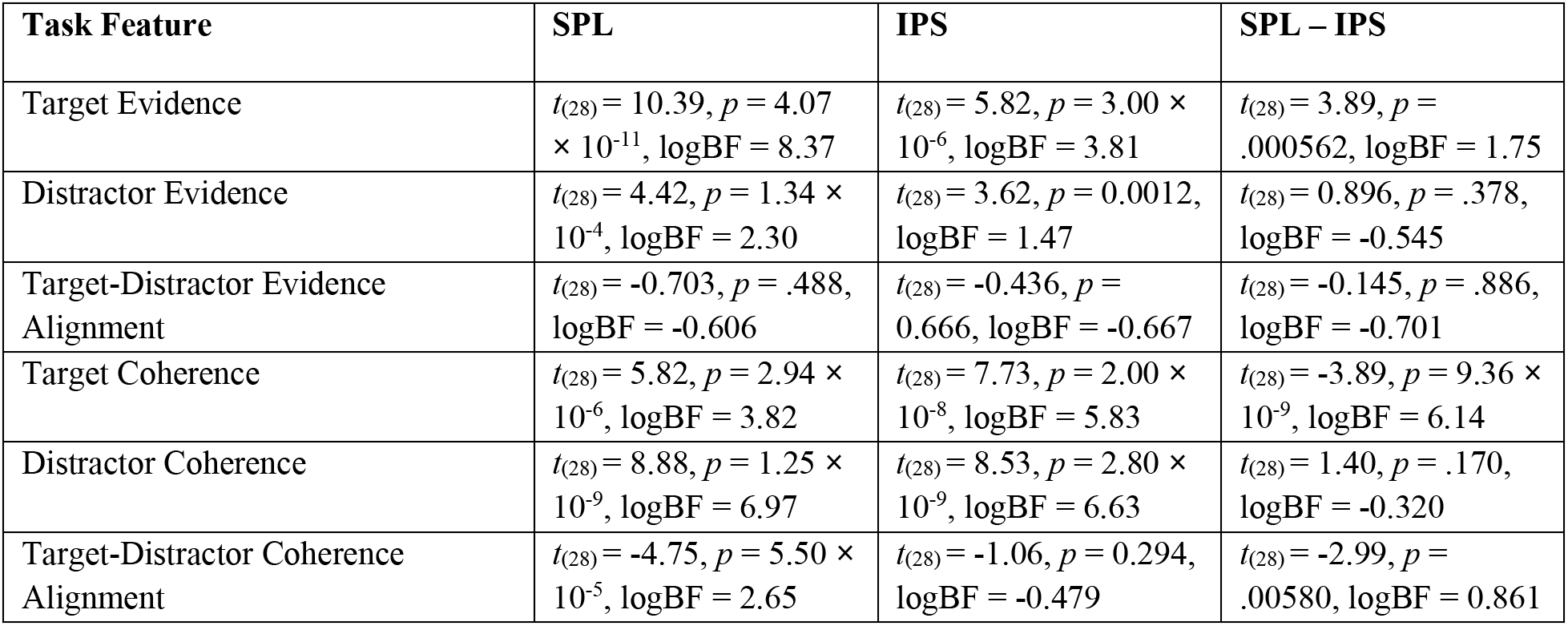
*Feature encoding contrasted across parietal cortex.* Encoding of feature evidence and coherence within SPL, within IPS, and contrasted between SPL and IPS. Note the stronger target evidence encoding in SPL, stronger target coherence encoding in IPS, and stronger target-distractor coherence alignment in SPL.

To further characterize how feature coherence and evidence are encoded across these regions, we performed multidimensional scaling over each regions task representations (Figure 4b; (Diedrichsen and Kriegeskorte, 2017; Kriegeskorte et al., 2006)). Briefly, this method allows us to visualize – in a non-parametric manner – the relationships between representations of different feature levels (e.g., levels of target coherence), by estimating each feature level separately within a GLM and then using singular value decomposition to project these patterns into a 2D space (see Methods for additional details). We found that coherence and evidence axes naturally emerge in the top two principal components in this analysis within dACC, SPL, and IPS. Coherence axes (light to dark shading) are parallel between left (blue) and right (brown) responses, suggesting a response-independent encoding. In these components, evidence encoding appeared to be binary, in contrast to parametric coherence encoding (we found similar whole-brain encoding maps for binary-coded evidence; see Supplementary Figure 7). Critically, whereas coherence encoding axes within SPL were aligned between targets (circles) and distractors (diamonds; confirming aligned encoding), in IPS these representations form perpendicular lines (confirming orthogonal encoding). In addition to visualizing cross-region dissociations in these representations, these analyses also help to validate that task features are encoded in a monotonic fashion.

Finally, to explore the divisions between SVA and Control networks evident in the univariate analyses, we split up our two prefrontal ROIs by their network membership (Supplementary Figure 8). In dACC, we found that SVA parcels tended to have stronger feature encoding than Control parcels. Interestingly, in these SVA parcels several features were aligned with the target evidence dimension, consistent with recent human electrophysiology findings (Ebitz et al., 2020). In lPFC, we found that Control parcels, but not SVA parcels, encoded distractor congruence (Control: *t*(28) = 3.60, two-tailed *p* = .0012, logBF = 1.45; SVA: *t*(28) = 0.57, *p* = .57, logBF = -0.64; Control – SVA: *t*(28) = 3.27, *p* = .0029, logBF = 1.12). This distractor congruence encoding was present in lPFC both in ‘Control A/B’ parcels (*t*(28) = 3.66, *p* = .001) and marginally in ‘Control C’ parcels (*t*(28) = 1.86, *p* = .073). This network-selective encoding of congruence is consistent with the univariate results in dACC (see Figure 2).

### Control demands dissociate coherence and evidence encoding

Our findings thus far demonstrate two sets of dissociations within and across brain regions. In dACC, we find that distinct regions encode the control demands related to discriminating targets (caudal dACC) versus overcoming distractor incongruence (rostral dACC). In posterior parietal cortex, we find that overlapping regions track the coherence of these two stimulus features, but that distinct regions represent these features in alignment (SPL) versus orthogonally (IPS). While these findings suggest that this set of regions was involved in translating between feature information and goal-directed responding, they only focus on the information that was presented to the participant on a given trial. To provide a more direct link between feature-specific encoding and control, we examined how the encoding of feature coherence differed between matched task that placed stronger or weaker demands on cognitive control. So far, our analyses have focused on conditions in which participants needed to respond to the color feature while ignoring the motion feature (Attend-Color task), but on alternating scanner runs participants instead responded to the motion dimension and ignored the color dimension (Attend-Motion task). These tasks were matched in their visual properties (identical stimuli) and motor outputs (left/right responses), but critically differed in their control demands. Attend-Motion was designed to be much easier than Attend-Color, as the left/right motion directions are compatible with the left/right response directions (i.e., Simon facilitation; (Danielmeier et al., 2011; Ritz and Shenhav, 2021)). Comparing these tasks allows us to disambiguate bottom-up attentional salience from the top-down contributions to attentional priority (Jackson et al., 2017; Woolgar et al., 2011a, 2015b, 2015a).

Consistent with previous work (Ritz and Shenhav, 2021), performance on the Attend-Motion task was better overall (mean RT: 565ms vs 725ms, sign-rank *p* = 2.56 **×** 10^-6^; mean Accuracy: 93.7% vs 87.5%, sign-rank *p* = .000318). Unlike the Attend-Color task, performance was not impaired by distractor incongruence (i.e., color distractors; RT: *t*(28) = -1.39, *p* = .176; Accuracy: *t*(28) = 0.674, *p* = .506). To investigate these task-dependent feature representations, we fit a GLM that included both tasks. To control for performance differences across tasks, we only analyzed accurate trials and included trial-wise RT as a nuisance covariate, concatenating RT across tasks.

Whereas the encoding of both color and motion coherence was widespread during the Attend-Color task (Figure 3), coherence encoding was consistently weaker during the less demanding Attend-Motion task (Figure 5A). Coherence encoding was weaker during Attend-Motion whether classifying according to goal-relevance (comparing targets or distractors) or the features themselves (comparing motion or color). Task-relevant ROIs revealed that coherence encoding was effectively absent during the easy Attend-Motion task (Figure 5B), suggesting that they depend on the control demands of the Attend-Color task (Woolgar et al., 2011a, 2011b).

**Figure 5.**
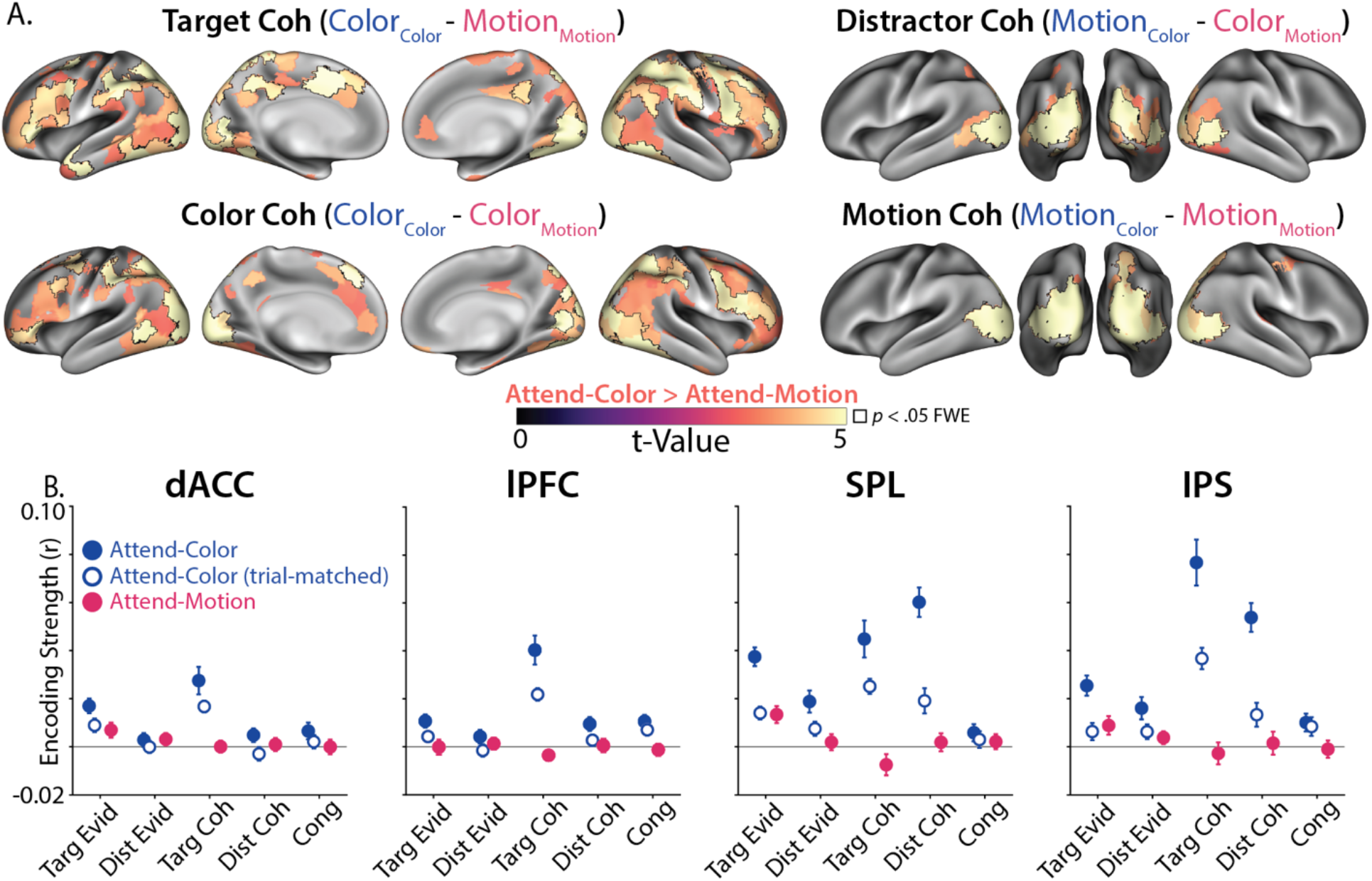
*Task-dependent encoding strength.* **A)** Across cortex, feature coherence encoding was stronger during Attend-Color than Attend-Motion, matched for the same number of trials. Attend-Color had stronger encoding when comparing target coherence (top left), distractor coherence (top right), color coherence (bottom left) and motion coherence (bottom right). Parcels are thresholded at *p* < .001 (uncorrected); outlined parcels are significant at *p* < .05 I (max-statistic randomization test across all parcels). Titles are coded ‘FeatureTask’. **B)** Target and distractor coherence information was encoded more strongly during Attend-Color than Attend-Motion in dACC, lPFC, SPL and IPS. Attend-Color encoding plotted from the whole sample (blue fill) and a trial-matched sample (first 45 trials of each run; white fill) In Attend-Motion runs, only target evidence was significantly encoded (magenta). **C)** Target and distractor coherence was not reliably encoded during the Attend-Motion task (liberally thresholded at *p* < .01 uncorrected). GLM: Between-Task, see Table 2.

In contrast to these stark task-related differences in coherence encoding, we found that neural encoding of the target evidence (color evidence in the Attend-Color task and motion evidence in the Attend-Motion task) was preserved across tasks, including within dACC, lPFC, SPL, and IPS (Figure 5B). Consistent with previous experiments examining context-dependent decision-making (Aoi et al., 2020; Flesch et al., 2022; Jackson et al., 2017; Kayser et al., 2010b; Mante et al., 2013; Pagan et al., 2022; Takagi et al., 2021), we found stronger target evidence encoding relative to distractor evidence encoding, in our case in the evidence-encoding SPL (Attend-Color: *t*(28) = 4.26, one-tailed *p* = 0.0001; Attend-Motion: *t*(28) = 2.37, one-tailed *p* = 0.0124). We also found that target evidence encoding during Attend-Motion was aligned with Attend-Color, both for *motion* evidence encoding (‘stimulus axis’; SPL: one-tailed *p* = .0236, IPS: one-tailed *p* = .0166) and *target* evidence encoding (‘decision axis’; SPL: one-tailed *p* = 1.29 **×** 10^-6^; IPS: one-tailed *p* = .0005), again in agreement with these previous experiments. Whereas our experiment replicates previous observations of the neural representations supporting contextual decision-making, we now extended these findings to understand how putative attention signals (i.e., feature coherence) are encoded in response to the asymmetric inference that is characteristic of cognitive control (Miller and Cohen, 2001).

### Aligned encoding dimensions for feature coherence and task performance

We next explored whether the encoding of feature coherence, seemingly in the service of cognitive control, was related to how well participants performed the task. We tested this question by determining whether feature coherence representations were aligned with representations of behavior (i.e., alignment between stimulus and behavioral subspaces (Stringer et al., 2019)). Specifically, we included trial-level reaction time and accuracy in our first-level GLMs. Encoding of performance was itself highly robust: most parcels encoded reaction time and accuracy, with the strongest encoding in cognitive control regions (Supplementary Figure 9). Across cortex, reaction time and accuracy were negatively correlated, again most prominently across the cognitive control network. To explore the behavioral relevance of coherence representations, we tested whether coherence encoding was aligned with the voxel patterns encoding task performance.

We found that the encoding of target and distractor coherence was aligned with performance across frontoparietal and visual regions (Figure 6a-b). If a regions’ encoding of target coherence reflects how sensitive the participant was to target information on that trial (e.g., due to top-down priority), we would expect target encoding to be positively aligned with performance on a given trial, such that stronger target coherence encoding is associated with better performance and weaker target coherence encoding is associated with poorer performance. We would also expect distractor encoding to demonstrate the opposite pattern – stronger encoding associated with poorer performance and weaker encoding associated with better performance. We found evidence for both patterns of feature-performance alignment across visual and frontoparietal cortex: target encoding was aligned with better performance (faster RTs and higher accuracy; Figure 6a), whereas distractor encoding was aligned with worse performance (slower RTs and lower accuracy; Figure 6b).

**Figure 6.**
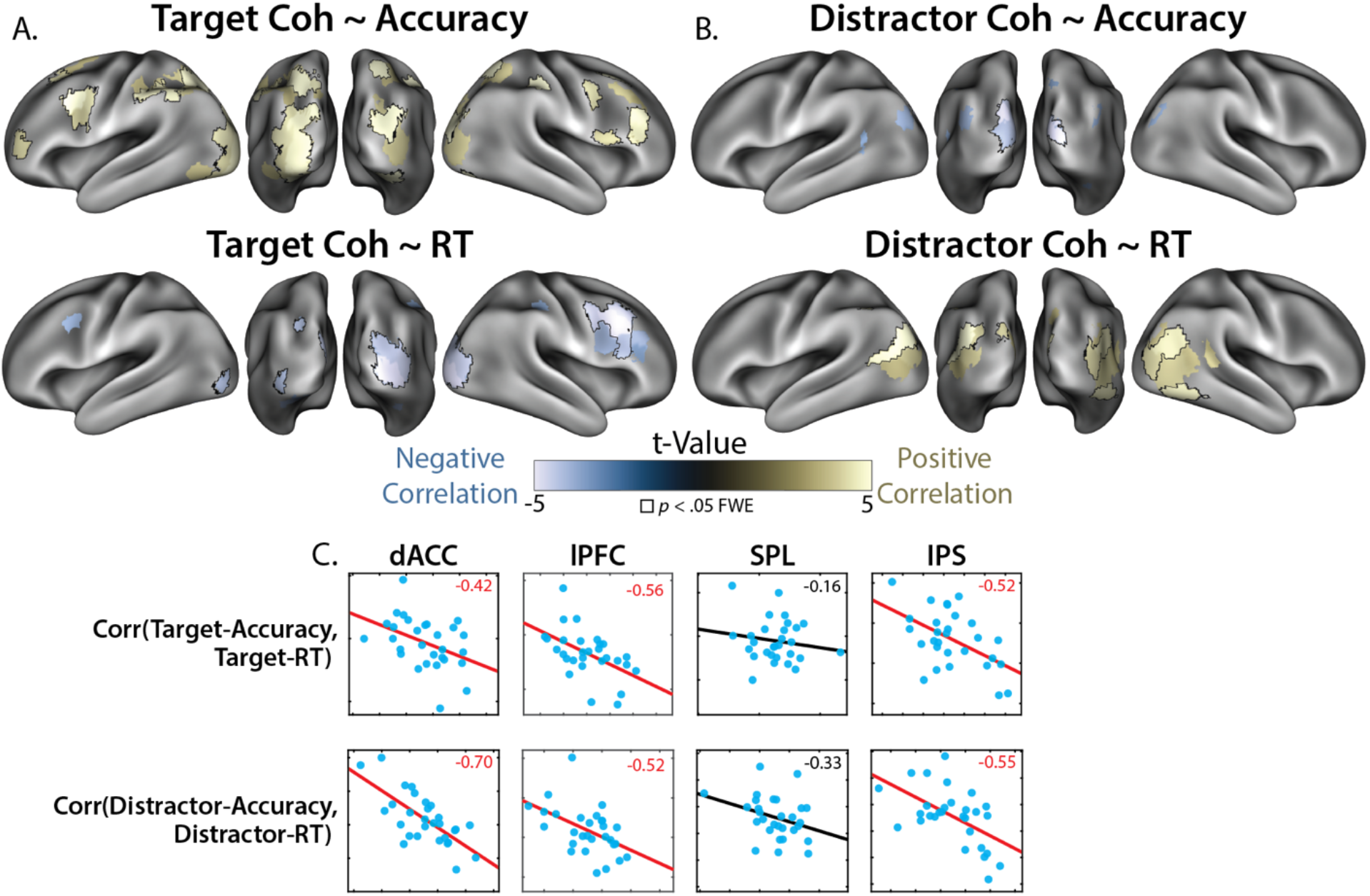
*Alignment between feature and performance encoding.* **A)** Alignment between encoding of target coherence and performance (top row: Accuracy, bottom row: RT). **B)** Alignment between encoding of distractor coherence and performance (top row: Accuracy, bottom row: RT). Across A and B, parcels are thresholded at *p* < .001 (uncorrected, in jointly reliable parcels), and outlined parcels are significant at *p* < .05 I (max-statistic randomization test across jointly reliable parcels). **C)** Individual differences in feature-RT alignment correlated with feature-accuracy alignment across regions (correlation values in top right; *p* < .05 in red). See Supplementary Table 1 for partial correlations controlling for reliability. GLM: Performance, see Table 2.

Next, we examined whether performance-coherence alignment reflected individual differences in participants’ task performance in our main task-related ROIs (see Figures 3-4). In particular, we tested whether the alignment between features and behavior reflects specific relationships with speed or accuracy, or whether they reflected overall increases in evidence accumulation (e.g., faster responding and higher accuracy). Within each ROI, we correlated feature-RT alignment with feature-accuracy alignment across subjects. We found that in dACC and IPS, participants showed the negative correlation between performance alignment measures predicted by an increase in processing speed (Figure 6c). People with stronger alignment between target coherence and shorter RTs tended to have stronger alignment between target coherence and higher accuracy, with the opposite found for distractors. While these between-participant correlations were present within targets and distractors, we did not find any significant correlations across features (between-feature: all *p*s > .10), again consistent with feature-specific processing. These analyses were qualitatively similar after partialing out the reliability of coherence and performance encoding, albeit with dACC and lPFC now showing marginal correlations for target coherence (see Supplementary Table 1). While between-participant analyses using small sample sizes warrant a note of caution, these findings are consistent across features and regions. In conjunction with our within-participant evidence that feature coherence representations are aligned with performance efficiency, these findings support a role for coherence encoding in adaptive control.

### Coherence encoding aligns with frontoparietal activity

Across frontal, parietal, and visual cortex, encoding of target and distractor coherence depended on task demands and was aligned with performance. Since this widespread encoding of task information likely reflects distributed network involvement in cognitive control (Corbetta and Shulman, 2002; Goldman-Rakic, 1988; Miller and Cohen, 2001), we sought to understand how frontal and parietal systems interact. We focused our analyses on IPS and lateral PFC (lPFC), linking the core parietal site of orthogonal coherence encoding (IPS) to an prefrontal site previous work suggests provides top-down feedback during cognitive control (Goldman-Rakic, 1988; Kastner and Ungerleider, 2000; Suzuki and Gottlieb, 2013; Yantis and Serences, 2003). Previous work has found that IPS attentional biases lower-level stimulus encoding in visual cortices (Kay and Yeatman, 2017; Saalmann et al., 2007), and that IPS mediates directed connectivity between lPFC and visual cortex during perceptual decision-making (Kayser et al., 2010b). Here, we extended these experiments to test how IPS mediates the relationship between prefrontal feedback and stimulus encoding.

To investigate these putative cortical interactions, we developed a novel multivariate connectivity analysis to test whether coherence encoding was aligned with prefrontal activity, and whether this lPFC-coherence alignment was mediated by IPS. We first estimated the voxel-averaged residual timeseries in lPFC (SPM12’s eigenvariate), and then included this residual timeseries alongside task predictors in a whole-brain regression analysis (Figure 7A). This analysis can be schematized as:

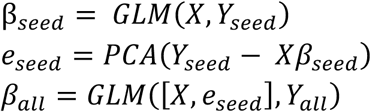

**Figure 7.**
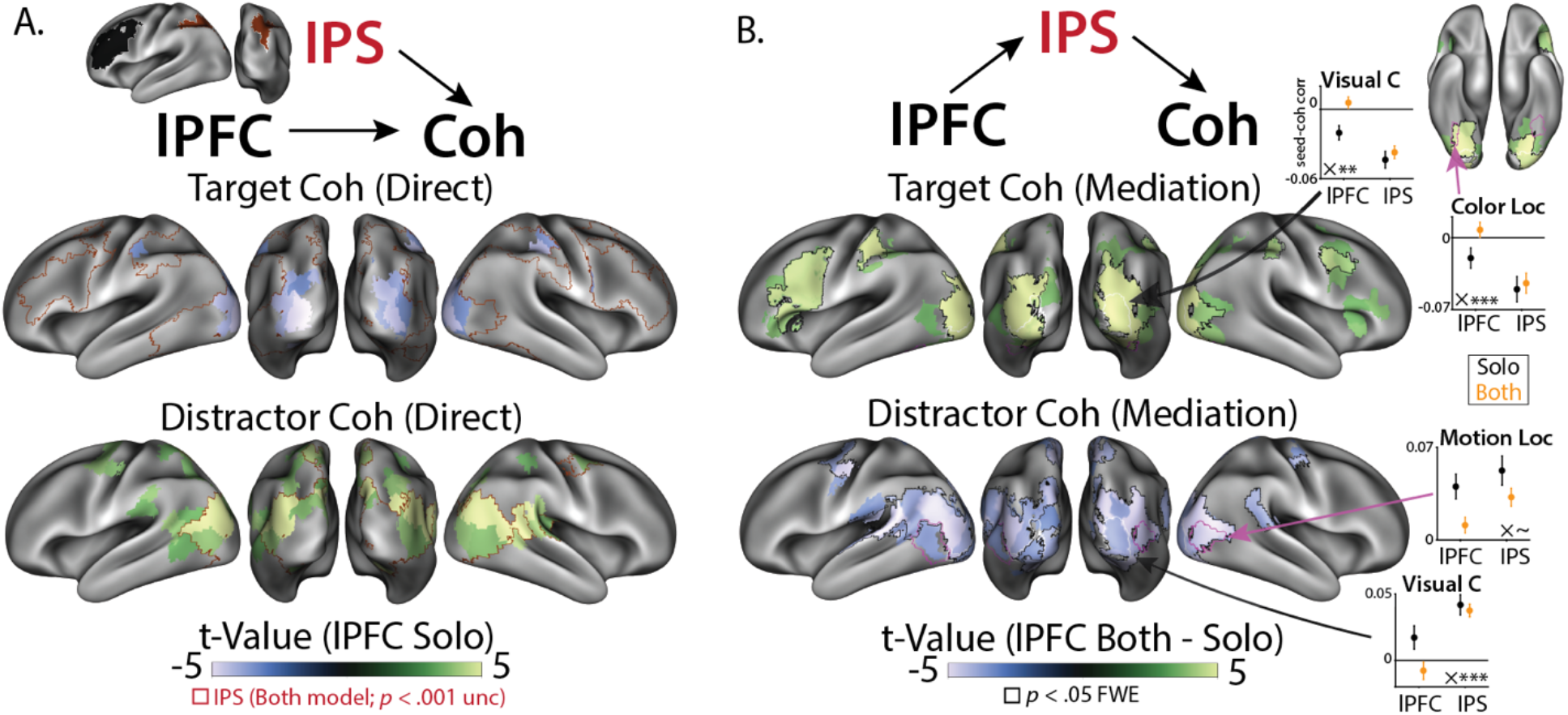
*IPS mediates alignment between lPFC and feature encoding.* **A)** Connectivity patterns from lPFC (color) and IPS (red outline) were aligned with target and distractor coherence patterns (*p* < .001 uncorrected, in jointly reliable parcels). IPS effects are outlined to show overlap, with all effects in a consistent direction to lPFC. **B)** lPFC-feature alignment contrasted between lPFC-only model (‘Solo’) and lPFC + IPS model (‘Both’). Including IPS reduced the alignment between lPFC and feature encoding (compare the sign of the main effect in B to the contrast in C). Parcels are thresholded at *p* < .001 (uncorrected, jointly reliable parcels), and outlined parcels are significant at *p* < .05 I (max-statistic randomization test across jointly reliable parcels). Insets graphs: seed-coherence alignment in Solo models (black) and Both model (orange) across visual regions. ‘Visual C’ is defined by our parcellation (Kong et al., 2021); Color and Motion localizers are parcels near the peak response identified during feature localizer runs (see Methods). In general, lPFC alignment was more affected by IPS than IPS alignment was affected by lPFC (inset ‘X’: difference of differences; ∼ *p* < .10, * *p* < .01, *** *p* < .001). GLM: Performance CX, see Table 2.

The GLM function performs regression using design matrix X and multivariate voxel timeseries Y, and the PCA function extracts the first principal component of the residuals. Finally, we used EGA to test whether there was alignment between patterns encoding lPFC functional connectivity (i.e., betas from the residual timeseries predictor e_seed) and patterns encoding target and distractor coherence. Note that while these results reflect functional connectivity, all correlational measures are subject to potential confounding (Reid et al., 2019).

We found that lPFC connectivity patterns were aligned with coherence-encoding patterns in visual cortex (Figure 7B). Stronger prefrontal functional connectivity was aligned with weaker target coherence and stronger distractor coherence, consistent with prefrontal recruitment during difficult trials. Notably, IPS connectivity was also aligned with target and distractor coherence in overlapping parcels, even when controlling for lPFC connectivity. These effects were liberally thresholded for visualization, as significant direct and indirect effects are not necessary for significant mediation (MacKinnon et al., 2007).

Our critical test was whether IPS mediated the relationship between lPFC activity and coherence encoding. We compared regression estimates between a model that only included lPFC residuals (‘solo’ model) to a model that included both lPFC and IPS residuals (‘both’ model). Comparing the strength of lPFC-coherence alignment with and without IPS is a test of whether parietal cortex mediates lPFC-coherence alignment (MacKinnon et al., 2007). These models can be schematized as:

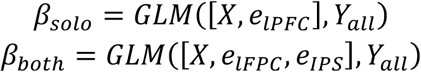

We found that this mediation was strongest in early visual cortex, where the alignment between lPFC and feature coherence was reduced in a model that included IPS relative to a model without IPS (Figure 7C). The negatively correlated target-lPFC relationship became more positive when IPS was included (top), and the positively correlated distractor-PFC relationship became more negative when IPS was included (bottom). Critically, we found that IPS reduced prefrontal-coherence alignment in early visual cortex more than lPFC reduced parietal-coherence alignment (Figure 7C inset; Supplementary Figure 10a-b), consistent with frontal-to-parietal directed connectivity in previous research (Kayser et al., 2010b; Suzuki and Gottlieb, 2013). Looking within color- and motion-sensitive parcels, determined using task-free localizer runs (see Methods), we found this mediation was robustly significant in color-sensitive cortex and marginally significant in motion-sensitive cortex. The opposite relationship, lPFC mediation of IPS connectivity, appeared in higher-level visual cortex for distractor coherence (Supplementary Figure 10c-d), though these effects were not reliable in explicit contrasts and may reflect projections from both regions.

While we were primarily interested in alignment with lPFC due to previous work implicating these regions in top-down control (for reviews, see (Friedman and Robbins, 2021; Shenhav et al., 2013)), for completeness we also examined how different subnetworks in both lPFC and dACC aligned with coherence encoding. In lPFC, we found that SVA and Control subnetworks had similar patterns of alignment (Supplementary Figure 11). In dACC we found that the SVA subnetwork had a qualitatively similar profile of coherence alignment as lPFC, but this alignment was absent in the Control subnetwork. Whereas this seed-coherence alignment was similar across lPFC and SVA dACC, unlike lPFC we found that SVA dACC failed to demonstrate strong evidence for mediation by IPS (Supplementary Figure 12).

A final set of analyses examined whether SPL and IPS demonstrated different patterns of task-related functional connectivity with other regions, given that we found that these regions differentially encoded evidence and coherence. When seeding our connectivity analyses with SPL activity, we found that SPL activity aligned with evidence encoding in bilateral motor cortex (Supplementary Figure 13). In contrast, IPS activity did not significantly align with evidence encoding, and this seed-evidence alignment in motor cortex was stronger for SPL than IPS, consistent with a putative role for SPL in response selection (Hunt et al., 2012).

## Discussion

In this experiment, we explored whether neural control systems use representations with the same dimensionality as the processes they regulate (Badre et al., 2021; Kalman, 1960; Ritz et al., 2022a). Inspired by behavioral evidence that participants can independently control their sensitivity to targets and distractors (Ritz and Shenhav, 2021), we set out to understand whether the neural correlates of monitoring and prioritization leverage independent encoding for feature-selective control (Figures 1a-c). We found that key nodes of canonical cognitive control networks had orthogonal neural representations of targets and distractors. Within dACC, orthogonal representations of target and distractor difficulty arose from segregated encoding along a rostrocaudal axis. Within IPS, orthogonal representations of target and distractor coherence arose from orthogonal subspaces in overlapping voxels. Consistent with a role in attentional priority, coherence representations depended on control demands, task performance, and frontoparietal activity. Together, these results reveal a neural mechanism for how cognitive control prioritizes multiple streams of information during decision-making.

Neurocomputational theories have proposed that dACC is involved in planning control across multiple levels of abstraction (Holroyd and McClure, 2015; Holroyd and Yeung, 2011; Shenhav et al., 2013; Vassena et al., 2017). Past work has found that control abstraction is hierarchically organized along dACC’s rostrocaudal axis, with more caudal dACC involved in lower-level action control, and more rostral dACC involved in higher-level strategy control (Shenhav et al., 2018; Taren et al., 2011; Venkatraman et al., 2009; Zarr and Brown, 2016), an organization that may reflect a more general hierarchy of abstraction within PFC (Badre and D’Esposito, 2009; Badre and Nee, 2018; Koechlin and Summerfield, 2007; Taren et al., 2011). Consistent with this account, we found that caudal dACC tracked the coherence of the target and distractor dimensions, especially within the SVA network. In contrast, more rostral dACC tracked incongruence between targets and distractors, especially within the Control network. Speculatively, our results are consistent with caudal dACC tracking the first-order difficulty arising from the relative salience of feature-specific information, and more rostral dACC tracking the second-order difficulty arising from cross-feature (in)compatibility (Badre and D’Esposito, 2009), the latter of which may require additional disengagement from distractor-dependent attentional capture.

Whereas dACC encoded feature difficulty (e.g., distractor incongruence), in parietal cortex we found overlapping representations of feature coherence (e.g., distractor coherence). In SPL, features had correlated coherence encoding (similarly representing low target coherence and high distractor coherence), consistent with this region’s transient and non-selective role in attentional control (Esterman et al., 2009; Greenberg et al., 2010; Molenberghs et al., 2007; Serences et al., 2004; Serences and Yantis, 2007; Yantis et al., 2002). In contrast, IPS had orthogonal representations of feature coherence, consistent with selective prioritization of task-relevant information (Adam and Serences, 2021; Greenberg et al., 2010; Jackson et al., 2017; Kay and Yeatman, 2017; Molenberghs et al., 2007; Serences and Yantis, 2007; Suzuki and Gottlieb, 2013; Woolgar et al., 2015b, 2015a, 2011a; Yantis et al., 2002). Our previous work has demonstrated behavioral evidence for independent control over target and distractor attentional priority in this task (Ritz and Shenhav, 2021), with different task variables selectively enhancing target or distractor sensitivity (see also (Egner, 2008; Soutschek et al., 2015)). Orthogonal feature representation in IPS may offer a mechanism for this feature-selective control, consistent with theoretical accounts of IPS implementing a priority map that combines stimulus- or value-dependent salience with goal-dependent feedback from PFC (Bisley and Goldberg, 2010; Bisley and Mirpour, 2019; Corbetta and Shulman, 2002; Gottlieb et al., 2020; Yantis and Serences, 2003).

In dACC, we found that target and distractor difficulty encoding was consistent with the segregated encoding hypothesis, with features evoking univariate responses in distinct but adjacent regions. Interestingly, we did not find corresponding encoding of distractor congruence in our multivariate analyses within dACC, potentially reflecting the spatial smoothness of this response. However, we did find multivariate encoding of distractor congruence in lPFC, and multivariate encoding of target and distractor coherence in IPS. These multivariate profiles were consistent with our subspace encoding hypothesis. The reasons for a mix of segregated and subspace encoding across cortex is unclear, but this may speculatively reflect the segregation across functional networks. Like in dACC, distractor congruence had stronger encoding within the lPFC Control network, albeit without the feature segregation (lPFC Control parcels also encoded target coherence in an orthogonal subspace). It is possible that these network segregations help bind related control processes (Corbetta and Shulman, 2002; Gordon et al., 2017; Menon and D’Esposito, 2021), a hypothesis that future experiments should test with targeted paradigms (e.g., with subject-specific functional networks).

By comparing two different task goals (Attend-Color vs. Attend-Motion), our study was able to test whether coherence representations reflect control-dependent prioritization of information processing. Previous research has shown that these tasks differ dramatically in their control demands (Ritz and Shenhav, 2021). As in previous work, task performance was much better in Attend-Motion runs than Attend-Color runs, and participants were not sensitive to color distractors. Consistent with previous work on context-dependent decision-making, target evidence had similarly strong encoding across tasks, with generalizable encoding dimensions for choice and motion directions (Flesch et al., 2022; Mante et al., 2013; Takagi et al., 2021). In contrast to these putative decision representations, we found that coherence representations disappeared in the easier Attend-Motion task. On its own, weaker encoding of color distractors in Attend-Motion could be explained by the weaker bottom-up salience of the color dimension. However, the stark drop in the encoding of target (motion) coherence in these blocks cannot be similarly accounted for – these differences in target coherence encoding showed the opposite relationship expected from salience: better encoding of low-salience color targets (hard Attend-Color task) and weaker encoding of high-salience motion targets (easy Attend-Motion task). Instead, this encoding profile is consistent with previous research finding that feature decoding is stronger for more difficult tasks (Rust and Cohen, 2022; Woolgar et al., 2015b, 2015a, 2011a) or when people are incentivized to use cognitive control (Etzel et al., 2016; Hall-McMaster et al., 2019).

Critically, stimuli and responses were matched across tasks, helping to rule out alternative accounts of coherence encoding based on ‘bottom-up’ stimulus salience, decision-making, or eye movements. Difficulty-dependent coherence encoding may instead reflect the involvement of an attention control system that can separately regulate target and distractor processing, speculatively indexing the top-down ‘gain’ or ‘priority’ on these features (Bisley and Goldberg, 2010; Gottlieb et al., 2020; Yantis and Serences, 2003). Supporting this account, coherence representations in cognitive control regions like IPS were aligned with performance representations, with target encoding strength aligned with better performance and distractor encoding strength aligned with poorer performance. Individual difference in feature-performance alignment was correlated across features, consistent with these representations reflecting the underlying processes (e.g., priority) that give rise to behavior, rather than performance monitoring or surprise (which would likely have the opposite relationship, e.g., high target coherence aligned with poorer performance).

Classic models of prefrontal involvement in cognitive control (Desimone and Duncan, 1995; Kastner and Ungerleider, 2000; Miller and Cohen, 2001) propose that prefrontal cortex biases information processing in sensory regions. In line with this macro-scale organization, we found that coherence encoding in visual cortex was related to functional connectivity with the frontoparietal network. In particular, coherence encoding in visual cortex was aligned with patterns of functional connectivity to lateral prefrontal cortex, and this feature-seed relationship was mediated by IPS. The results of this novel multivariate connectivity analysis are consistent with previous research supporting a role for IPS in top-down control of visual encoding (Kay and Yeatman, 2017; Lauritzen et al., 2009; Saalmann et al., 2007), as well as a granger-causal PFC-IPS-visual pathway during a similar decision-making task (Kayser et al., 2010b). Here, we demonstrate stable ‘communication subspaces’ between visual cortex and PFC (Semedo et al., 2019; Srinath et al., 2021), which can plausibly communicate feedback adjustments to feature gain. With that said, while our interpretation of the direction of communication is therefore supported by prior work, these connectivity methods are correlational (Reid et al., 2019), and cannot rule out the possibility that our mediation findings reflect a bottom-up pattern of communication (e.g., visual-IPS-PFC). The asymmetric mediation between regions (i.e., IPS mediates lPFC more than lPFC mediates IPS; Supplementary Figure 10) rules out a range of potential confounders, and these regions were selected based on the anatomical connectivity within the frontoparietal network, notably through the superior longitudinal fasciculus (Petrides and Pandya, 2006). Future research should use temporally precise neuroimaging to account for directionality, causal manipulations to account for causality (e.g., (Jackson et al., 2021)), and should explore the higher dimensional connectivity subspaces that link different regions (Rust and Cohen, 2022; Srinath et al., 2021). These considerations notwithstanding, our findings are consistent with IPS, a critical site for orthogonal feature representations, playing a key role in linking prefrontal cortex with early perceptual processing.

Collectively, our findings provide new insights into how the brain may control multiple streams of information processing. While evidence for multivariate control has a long history in attentional tracking (Pylyshyn and Storm, 1988; Vul et al., 2009), including parametric relationships between attentional load and IPS activity (Culham et al., 2001, 1998; Howe et al., 2009; Jovicich et al., 2001; Ritz et al., 2022b), little is known about how the brain coordinates multiple control signals (Badre et al., 2021; Ritz et al., 2022a). Future experiments should further elaborate on this frontoparietal control circuit, for instance by interrogating how incentives influence different task representations (Etzel et al., 2016; Hall-McMaster et al., 2019; Parro et al., 2018; Peck et al., 2009; Wisniewski et al., 2015), or how neural and behavioral indices of control causally depend on perturbations of neural activity (Jackson et al., 2021). Future experiments should also use fast timescale neural recording technologies like (i)EEG or (OP-)MEG to better understand the within-trial dynamics of multivariate control (Ritz and Shenhav, 2021; Weichart et al., 2020). In sum, this experiment provides new insights into the large-scale neural networks involved in multivariate cognitive control, and points towards new avenues for developing a richer understanding of goal-directed attention.

## Methods

### Participants

Twenty-nine individuals (17 females, Age: M = 21.2, SD = 3.4) participated in this experiment. All participants had self-reported normal color vision and no history of neurological disorders. Two participants missed one Attend-Color block (see below) due to a scanner removal, and one participant missed a motion localizer due to a technical failure, but all participants were retained for analysis. Participants provided informed consent, in accordance with Brown University’s institutional review board.

### Task

The main task closely followed our previously reported behavioral experiment (Ritz and Shenhav, 2021). On each trial, participants saw a random dot kinematogram (RDK) against a black background. This RDK consisted of colored dots that moved left or right, and participants responded to the stimulus with button presses using their left or right thumbs.

In Attend-Color blocks (six blocks of 150 trials), participants responded depending on which color was in the majority. Two colors were mapped to each response (four colors total), and dots were a mixture of one color from each possible response. Dots colors were approximately isolument (uncalibrated; RGB: [239, 143, 143], [191, 239, 143], [143, 239, 239], [191, 143, 239]), and we counterbalanced their assignment to responses across participants.

In Attend-Motion blocks (six blocks of 45 trials), participants responded based on the dot motion instead of the dot color. Dot motion consisted of a mixture between dots moving coherently (either left or right) and dots moving in a random direction. Attend-Motion blocks were shorter because they acted to reinforce motion sensitivity and provide a test of stimulus-dependent effects.

Critically, dots always had color and motion, and we varied the strength of color coherence (percentage of dots in the majority) and motion coherence (percentage of dots moving coherently) across trials. Our previous experiments have found that in Attend-Color blocks, participants are still influenced by motion information, introducing a response conflict when color and motion are associated with different responses (Ritz and Shenhav, 2021). Target coherence (e.g., color coherence during Attend-Color) was linearly spaced between 65% and 95% with 5 levels, and distractor congruence (signed coherence relative to the target response) was linearly spaced between -95% and 95% with 5 levels. In order to increase the salience of the motion dimension relative to the color dimension, the display was large (∼10 degrees of visual angle) and dots moved quickly (∼10 degrees of visual angle per second).

Participants had 1.5 seconds from the onset of the stimulus to make their response, and the RDK stayed on the screen for this full duration to avoid confusing reaction time and visual stimulation (the fixation cross changed from white to gray to register the response). The inter-trial interval was uniformly sampled from 1.0, 1.5, or 2.0 seconds. This ITI was relatively short in order to maximize the behavioral effect, and because efficiency simulations showed that it increased power to detect parametric effects of target and distractor coherence (e.g., relative to a more standard 5 second ITI). The fixation cross changed from gray to white for the last 0.5 seconds before the stimulus to provide an alerting cue.

### Procedure

Before the scanning session, participants provided consent and practiced the task in a mock MRI scanner. First, participants learned to associate four colors with two button presses (two colors for each response). After being instructed on the color-button mappings, participants practiced the task with feedback (correct, error, or 1.5 second time-out). Errors or time-out feedback were accompanied with a diagram of the color-button mappings. Participants performed 50 trials with full color coherence, and then 50 trials with variable color coherence, all with 0% motion coherence. Next, participants practiced the motion task. After being shown the motion mappings, participants performed 50 trials with full motion coherence, and then 50 trials with variable motion coherence, all with 0% color coherence. Finally, participants practiced 20 trials of the Attend-Color task and 20 trials of Attend-Motion tasks with variable color and motion coherence (same as scanner task).

Following the twelve blocks of the scanner task, participants underwent localizers for color and motion, based on the tasks used in our previous experiments (Shenhav et al., 2018). Both localizers were block designs, alternating between 16 seconds of feature present and 16 seconds of feature absent for seven cycles. For the color localizer, participants saw an aperture the same size as the task, either filled with colored squares that were resampled every second during stimulus-on (‘Mondrian stimulus’), or luminance-matched gray squares that were similarly resampled during stimulus-off. For the motion localizer, participants saw white dots that were moving with full coherence in a different direction every second during stimulus-on, or still dots for stimulus-off. No responses were required during the localizers.

### MRI sequence

We scanned participants with a Siemens Prisma 3T MR system. We used the following sequence parameters for our functional runs: field of view (FOV) = 211 mm × 211 mm (60 slices), voxel size = 2.4 mm^3^, repetition time (TR) = 1.2 sec with interleaved multiband acquisitions (acceleration factor 4), echo time (TE) = 33 ms, and flip angle (FA) = 62°. Slices were acquired anterior to posterior, with an auto-aligned slice orientation tilted 15° relative to the AC/PC plane. At the start of the imaging session, we collected a high-resolution structural MPRAGE with the following sequence parameters: FOV = 205 mm × 205 mm (192 slices), voxel size = 0.8 mm^3^, TR = 2.4 sec, TE1 = 1.86 ms, TE2 = 3.78 ms, TE3 = 5.7 ms, TE4 = 7.62, and FA = 7°. At the end of the scan, we collected a field map for susceptibility distortion correction (TR = 588ms, TE1 = 4.92 ms, TE2 = 7.38 ms, FA = 60°).

### fMRI preprocessing

We preprocessed our structural and functional data using fMRIprep (v20.2.6; (Esteban et al., 2019) based on the Nipype platform (Gorgolewski et al., 2011). We used FreeSurfer and ANTs to nonlinearly register structural T1w images to the MNI152NLin6Asym template (resampling to 2mm). To preprocess functional T2w images, we applied susceptibility distortion correction using fMRIprep, co-registered our functional images to our T1w images using FreeSurfer, and slice-time corrected to the midpoint of the acquisition using AFNI. We then registered our images into MNI152NLin6Asym space using the transformation that ANTs computed for the T1w images, resampling our functional images in a single step. For univariate analyses, we smoothed our functional images using a Gaussian kernel (8mm FWHM, as dACC responses often have a large spatial extent). For multivariate analyses, we worked in the unsmoothed template space (see below).

### fMRI univariate analyses

We used SPM12 (v7771) for our univariate general linear model (GLM) analyses. Due to high trial-to-trial collinearity from to our short ITIs, we performed all analyses across trials, rather than extracting single-trial estimates. Our regression models used whole trials as events (i.e., a 1.5 second boxcar aligned to the stimulus onset). We parametrically modulated these events with standardized trial-level predictors (e.g., linear-coded target coherence, or contrast-coded errors), and then convolved these predictors with SPM’s canonical HRF, concatenating our voxel timeseries across runs. We included nuisance regressors to capture 1) run intercepts and 2) the average timeseries across white matter and CSF (as segmented by fMIRPrep). To reduce the influence of motion artifacts, we used robust weighted least-squares (Diedrichsen and Shadmehr, 2005; Jones et al., 2021), a procedure for optimally down-weighting noisy TRs.

We estimated contrast maps at the subject-level, which we then used for one-sample t-tests at the group-level. We controlled for family-wise error rate using threshold-free cluster enhancement (Smith and Nichols, 2009), testing whether voxels have an unlikely degree of clustering under a randomized null distribution (Implemented in PALM (Winkler et al., 2014); 10,000 randomizations). To improve the specificity of our coverage (e.g., reducing white-matter contributions) and to facilitate our inference about functional networks (see below), we limited these analyses to voxels within the Kong2022 whole-brain parcellation (Kong et al., 2021; Schaefer et al., 2018). This parcellation assigns regional labels to parcels (e.g., whether parcels are in ‘SPL’ or ‘IPS’), which was used through-out to generate ROIs. Surface renders were generated using surfplot (Gale et al., 2021; Vos de Wael et al., 2020), projecting from MNI space to the Human Connectome Project’s fsLR space (164,000 vertices).

### dACC longitudinal axis analyses

To characterize the spatial organization of target difficulty and distractor congruence signals in dACC, we constructed an analysis mask that provided broad coverage across cingulate cortex and preSMA. This mask was constructed by 1) getting a meta-analytic mask of cingulate responses co-occurring with ‘cognitive control’ (Neurosynth uniformity test; (Yarkoni et al., 2011), and taking any parcels from the whole-brain Schaefer parcellation (400 parcels; (Kong et al., 2021; Schaefer et al., 2018) that had a 50 voxel overlap with this meta-analytic mask. We used this parcellation because it provided more selective gray matter coverage than the Neurosynth mask alone and it allowed us to categorize voxels membership in putative functional networks.

To characterize the spatial organization within dACC, we first performed PCA on the masked voxel coordinates (y and z), getting a score for eac’ voxel’s position on the longitudinal axis of this ROI. We then regressed voxel’s gradient scores against their regression weights from a model including linear target coherence and distractor congruence (both coded -1 to 1 across difficulty levels). We used linear mixed effects analysis to partially pool across subjects and accommodate within-subject correlations between voxels. Our model predicted gradient score from the linear and quadratic expansions of the target and distractor betas (gradientScore ∼ 1 + target + target^2^ + distractor + distractor^2^ + (1 + target + target^2^ + distractor + distractor^2^ | subject)). To characterize the network-dependent organization of target and distractor encoding, we complexity-penalized fits between models that either 1) predicted target or distractor betas from linear and quadratic expansions of gradient scores, or 2) predicted target/distractor betas from dummy-coded network assignment from the Schaefer parcellation, comparing these models against a model that used both network and gradient information.

### Encoding Geometry Analysis (EGA)

We adapted functions from the pcm-toolbox and rsatoolbox packages for our multivariate analyses (Diedrichsen et al., 2018; Nili et al., 2014). We first fit whole-brain GLMs without spatial smoothing, separately for each scanner run. These GLMs estimated the parametric relationship between task variables and BOLD response (e.g., linearly coded target coherence), with a pattern of these parametric betas across voxels reflecting linear encoding subspace (Kriegeskorte and Diedrichsen, 2019). Within each Schaefer parcel (n=400), we spatially pre-whitened these beta maps, reducing noise correlations between voxels that can inflate pattern similarity and reduce reliability (Walther et al., 2016). We then computed the cross-validated Pearson correlation, estimating the similarity of whitened patterns across scanner runs. We used a correlation metric to estimate the alignment between encoding subspaces, rather than distances between condition patterns, to normalize biases and scaling across stimuli (e.g., greater sensitivity to targets vs distractors) and across time (e.g., representational drift). We found convergent results when using (un-centered) cosine similarity, suggesting that our results were not trivially due to parcels’ univariate response, but a correlation metric had the best reliability across runs. Note that this analysis approach is related to ‘Parallelism Scores’ (Bernardi et al., 2020), but here we use parametric encoding models and emphasize not only deviations from parallel/orthogonal, but also the direction of alignment between features (e.g., Figures 5 and 7).

We computed subspace alignment between contrasts of interest within each participant, and then tested these against zero at the group level. Since our correlations were less than *r* = |0.5|, we did not transform correlations before analysis. We used a Bayesian t-test to test for orthogonality (bayesFactor toolbox in MATLAB, based on (Rouder et al., 2012)). The Bayes factor from this t-test gives evidence for either non-orthogonality (BF10 further from zero) or orthogonality (BF10 closer to zero, often defined as the reciprocal BF01). Using a standard prior (Cauchy, width = 0.707), our strongest possible evidence for the orthogonality is BF01 = 5.07 or equivalently logBF = -0.705 (i.e., the Bayes factor when *t(28)* = 0).

Our measure of encoding strength was whether encoding subspaces were reliable across blocks (i.e., whether within-feature encoding pattern correlations across runs were significantly above zero at the group level). We used pattern reliability as a geometric proxy for how well a linear encoder would predict held-out brain data, as reliability provides the similarity between the cross-validated model and the best linear unbiased estimator of the within-sample data. We confirmed through simulations that pattern reliability is a good proxy for the traditional encoding metric of predicting held-out timeseries (Kriegeskorte and Diedrichsen, 2019). However, we found that pattern reliability is more powerful, due to it being much less sensitive to the magnitude of residual variance (these two methods are identical in the noise-free case; see Supplementary Figure 3).

When looking at alignment between two subspaces across parcels, we first selected parcels that significantly encoded both factors (‘jointly reliable parcels’, both *p* < .001 uncorrected). This selection process acts as a thresholded version of classical correlation disattenuation (Spearman, 1987; Thornton and Mitchell, 2017), and we confirmed through simulations this selection procedure does not increase type 1 error rate. We corrected for multiple comparisons using non-parametric max-statistic randomization tests across parcels (Nichols and Holmes, 2002). These randomization tests determine the likelihood of an observed effects under a null distribution generated by randomizing the sign of alignment correlations across participants and parcels 10,000 times. Within each randomization, we saved the max and min group-level effect sizes across all parcels, estimating the strongest parcel-wise effect we’d expect if there wasn’t a systematic group-level effect.

Some of our first-level models had non-zero levels of multicollinearity, due to conditioning on trials without omission errors or when including feature coherence and performance in the same model. Multicollinearity was far below standard thresholds for concern, generally (much) less than 5 for a standard threshold is 30 (ratio between largest and smallest singular values in the design matrix, using MATLAB colintest; (Belsley et al., 1980)). However, we wanted to confirm that predictor correlations wouldn’t bias our estimates of encoding alignment. We simulated data from a pattern component model (Diedrichsen et al., 2018) in which two variables were orthogonal (generated by separate variance components with no covariance), but were generated from a design matrix with correlated predictors. These simulations confirmed that cross-validated similarity measures were not biased by predictor correlations (Supplementary Figure 14).

To provide further validation for our parametric analyses, we estimated encoding profiles using an analysis with fewer parametric assumptions. First, we fit a GLM with separate predictors for levels of target and distractor evidence (‘Evidence Levels’ GLM in Table 2). Next, we estimated a traditional (cross-validated) representational dissimilarity matrix across all feature levels. Finally, we visualized these encoding profiles using classical multidimensional scaling (eigenvalue decomposition; see Figure 4B).

### Multivariate Connectivity Analysis

To estimate what information is plausibly communicated between cortical areas, we measured the alignment between multivariate connectivity patterns (i.e., the ‘communication subspace’ with a seed region, (Semedo et al., 2019)) and local feature encoding patterns. First, we residualized our Performance GLM (see Table 2) from a seed region’s timeseries, and then extracted the variance-weighted average timecourse (i.e., the leading eigenvariate from SPM12’s volume of interest function). We then re-estimated our Performance GLM, including the block-specific seed timeseries as a covariate, and performed EGA between seed and coherence patterns. We found convergent results when we residualized a quadratic expansion of our Performance GLM from our seed region, helping to confirm that connectivity alignment wasn’t due to underfitting. Note that our cross-validated EGA helps avoid false positives due to any correlations in the design matrix (see above). We localized this connectivity analysis to color- and motion-sensitive cortex by finding the bilateral Kong22 parcels that roughly covered the area of strongest block-level contrast during our localizer runs. Note that these analyses reflect ‘functional connectivity’, which is susceptible to unmodelled confounders (Reid et al., 2019).

To estimate the mediation of lPFC connectivity by IPS, we compared models in which just lPFC or just IPS were used for EGA against a model where both seeds were included as covariates in the same model (MacKinnon et al., 2007). Our test of mediation was the group-level difference in lPFC seed-coherence alignment before and after including IPS. While these analyses are inherently cross-sectional (i.e., lPFC and IPS are measured at the same time), we supplemented these analyses by showing that the mediating effect of IPS on lPFC was much larger than the mediating effect of lPFC on IPS (see Figure 7c; Supplementary Figure 10).

## Acknowledgements

This work was supported by NIH grant R01MH124849 (A.S.), NSF CAREER Award 2046111 (A.S.), NIH grant S10OD025181, and the C.V. Starr Postdoctoral Fellowship (H.R.). We are grateful to Joonhwa Kim for her assistance in data collection, and to Michael J. Frank, Matthew N. Nassar, Jonathan Cohen, Michael Esterman, Romy Frömer, Jörn Diedrichsen, Apoorva Bhandari, Debbie Yee, Sam Nastase, and the rest of the Shenhav Lab for helpful discussions.

## Data Availability

Data and analysis scripts will be made available upon publication.

## Supplementary Figures

**Figure S1.**
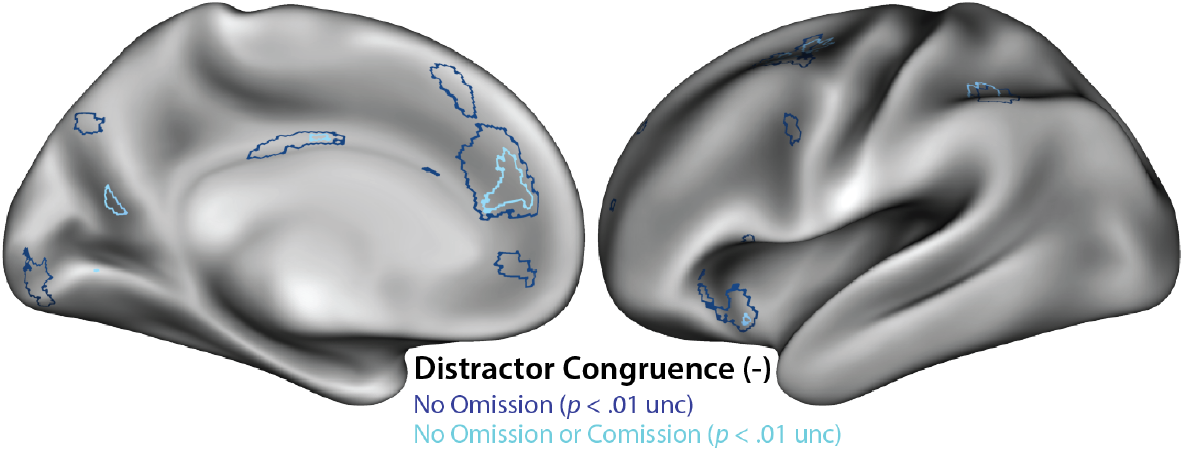
*Error control analysis.* Distractor congruence effect when controlling for different types of errors. Our primary analysis only analyzed trials without omission errors (navy), here plotted at a liberal uncorrected threshold. When we analyze trials without omission errors and commission errors (cyan), we see a consistent whole-brain topography, albeit at a lower statistical threshold. In both cases, relevant errors trials were included as nuisance events.

**Figure S2.**
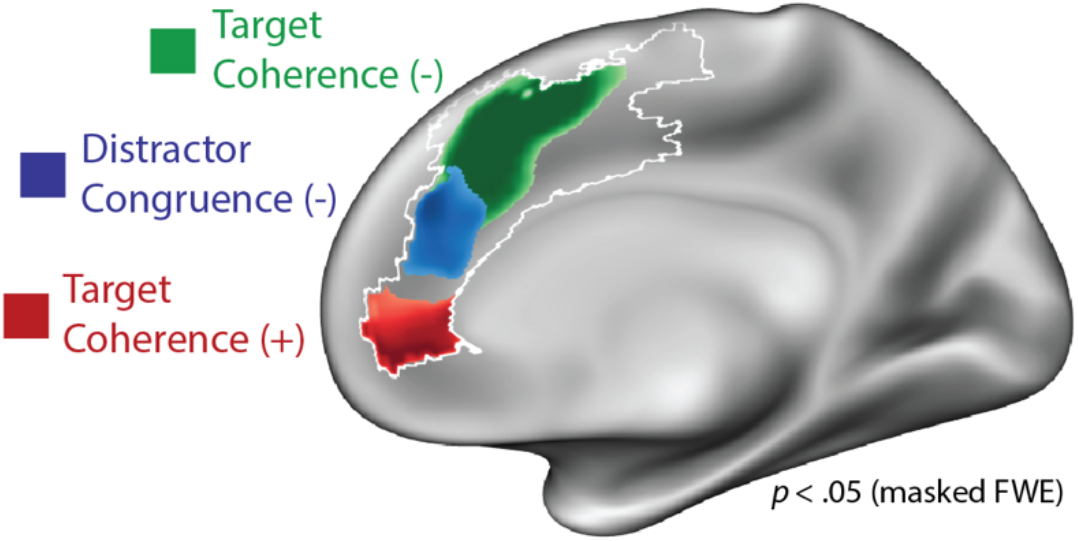
*Target ease.* Parametric effects of target coherence and distractor congruence, showing the rostral effect of target ease (positive relationship with target coherence) in red.

**Figure S3.**
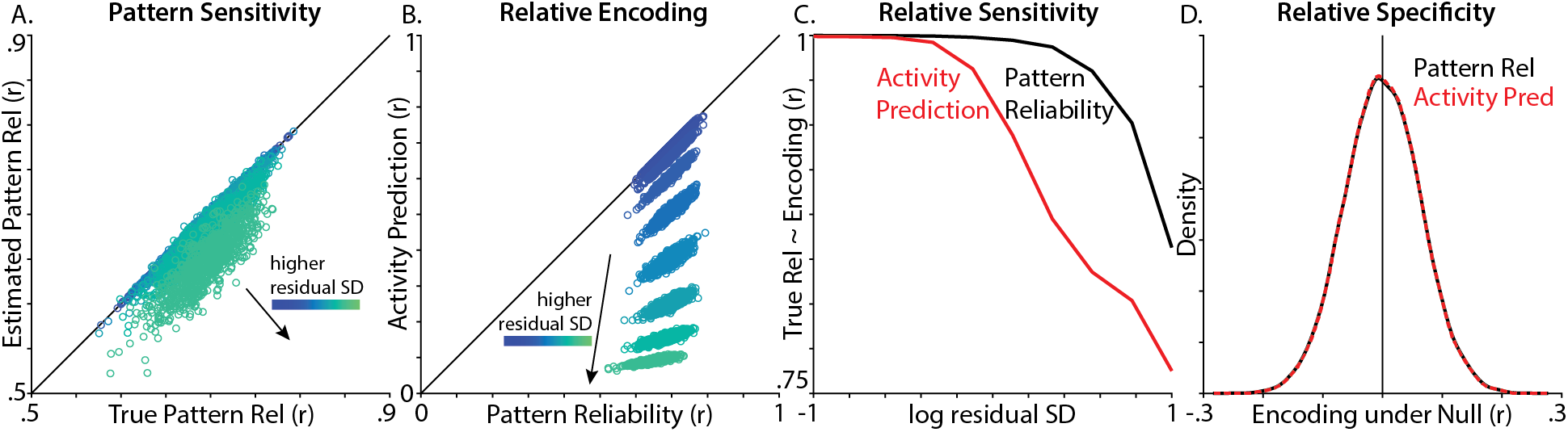
*Encoding Geometry Analysis (EGA) validation.* We validated how well we could recover the similarity between linear Gaussian models (training: *Y* = *XB* + Σ, test: *Y*′ = *X*′*B*′ + Σ). *Y* is the [1000 × 250] activity timeseries, *X* is the [1000 × 1] design matrix, *B* is the [1 × 250] encoding profile, and Σ reflects IID Gaussian noise. In each of our 1000 simulations, we used two different methods to recover the similarity between the true training encoding profile (*B*) and the true test encoding profile (*B*′ = *B* + 𝒩(0,1)), based on noisy activity timeseries (*Y* = *XB* + 𝒩(0, *σ_Y_*); *Y*′ = *X*′*B*′ + 𝒩(0, *σ_Y_*)). The first method was *pattern reliability* (i.e., our EGA method), correlating the encoding profile estimated during training 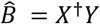, † indicates pseudoinverse) with the encoding profile estimated during test 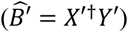. The second method was *activity prediction* (i.e., the traditional encoding approach), correlating the ground-truth test activity (*Y*′) with the predicted test activity 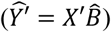. To simulate the high measurement noise inherent to fMRI, we compared these methods under different levels of residual SD (*σ_Y_*). **A)** Estimated pattern reliability tracked the true pattern reliability across the full range of residual SD, with some attenuation at high levels of noise **B)** Unlike pattern reliability, activity prediction became much poorer as residual SD increased. **C)** Correlating the true pattern reliability (correlation between *B* and *B*′) and estimated encoding strength (i.e., pattern reliability or activity prediction), we found pattern reliability was better correlated with the true reliability, particularly at higher levels of noise. **D)** Both methods had similar performance in the absence of a signal 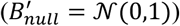).

**Figure S4.**
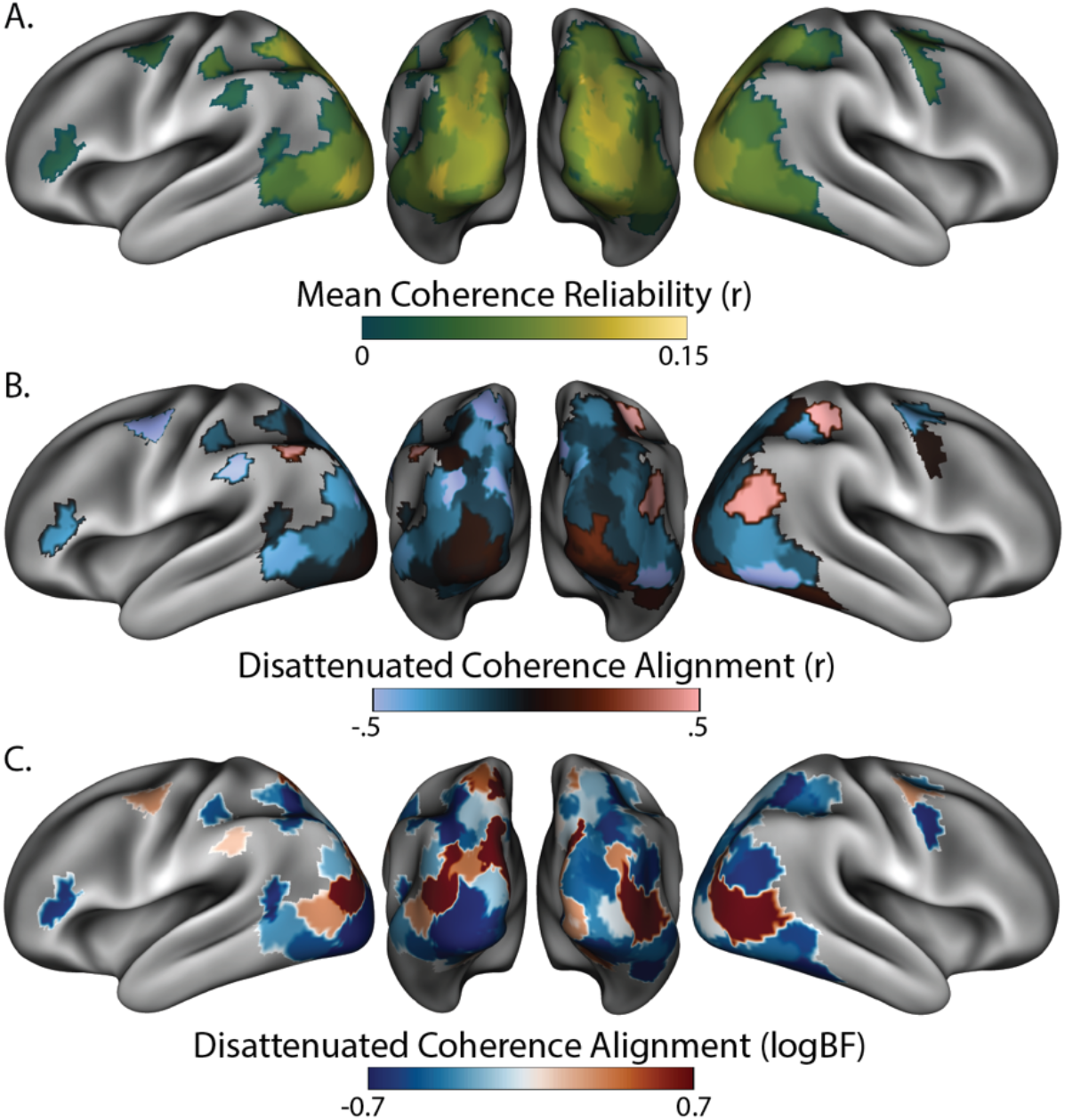
*Reliability control analysis.* **A)** Geometric mean of target and distractor coherence reliability 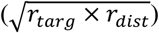, plotted in the reliability-thresholded parcels used in Figure 4. Reliability provides the theoretical upper bound on correlation strength. Median across participants, excluding participants with non-positive reliability. **B)** Target-distractor correlations, normalized by target-distractor reliability (i.e., disattenuated correlations) **C)** Log bayes factors for disattenuated target-distractor correlations. Compare to Figure 4C.

**Figure S5.**
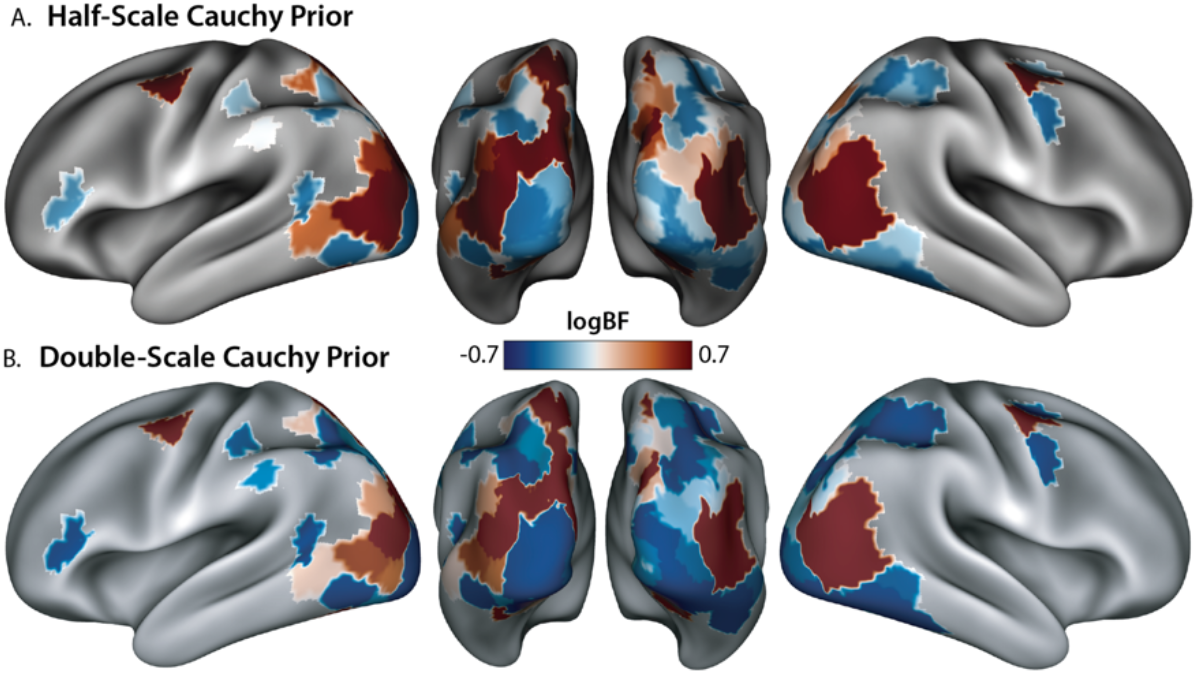
*Bayes factor prior control analysis.* **A)** Log bayes factors for target-distractor coherence alignment using a narrower prior (one-half the default Cauchy scale = 0.35). Minimum logBF is -0.46 at *t*(28) = 0. **B)** Same log bayes factor using a wider prior (double the default Cauchy scale = 1.41). Minimum logBF = -0.99 at *t*(28) = 0. Across different prior parameterizations, note the similarity to Figure 4C.

**Figure S6.**
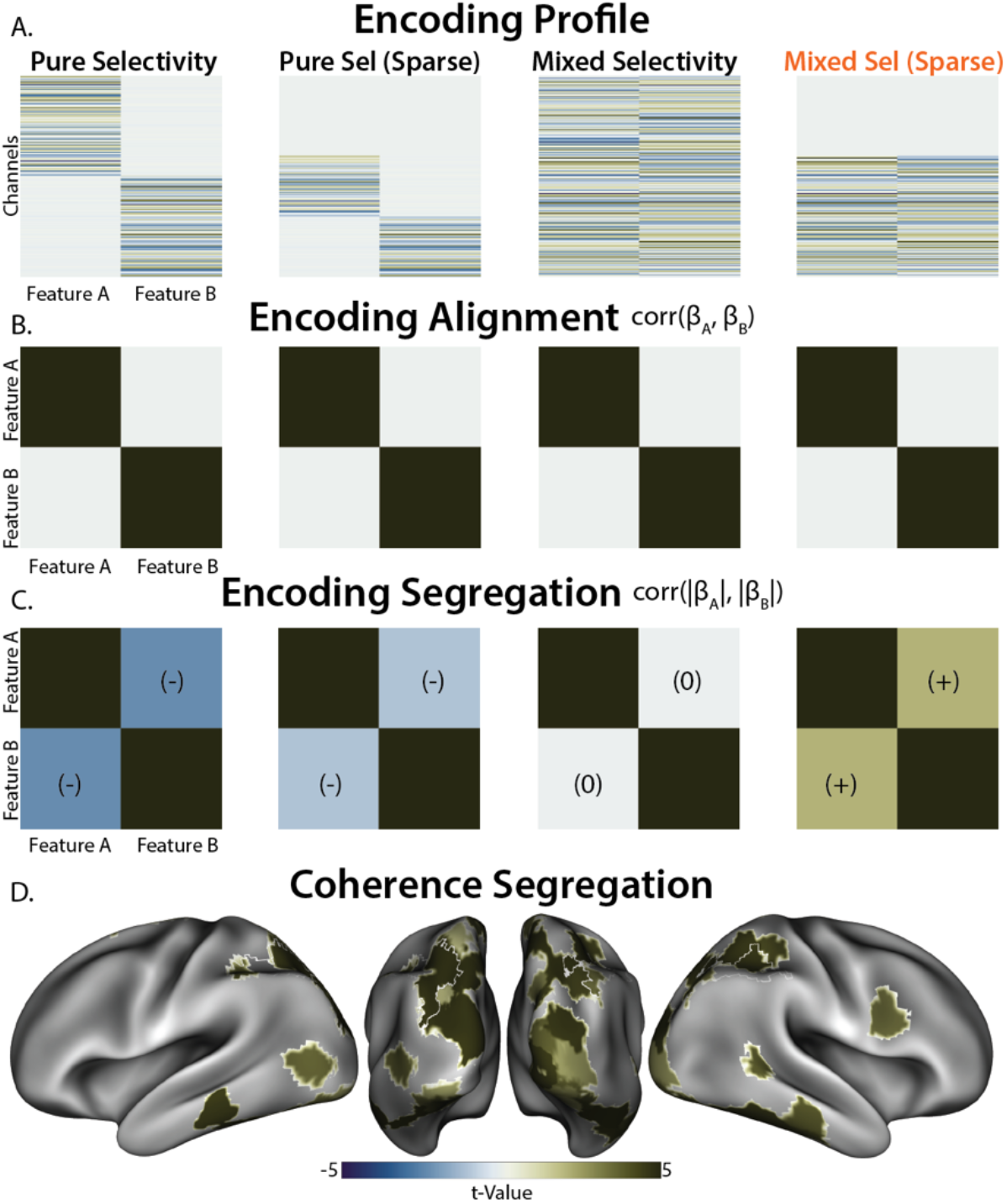
*Segregation Analysis.* **A)** We used pattern component modelling (Diedrichsen et al., 2018) to simulate different candidate encoding profiles. ‘Pure Selectivity’ reflects the segregated encoding hypothesis, with different voxels (rows) encoding different features (columns). ‘Mixed Selectivity’ reflects the orthogonal subspace hypothesis, with the same voxels encoding both features. ‘Sparse’ models include non-selective voxels. **B)** By design, all of these encoding profiles had the same orthogonal encoding alignment (uncorrelated encoding weights), highlighting that this measure is unable to adjudicate between candidate encoding profiles. **C)** These models can be differentiated by correlating their absolute encoding weights, testing whether the sensitivity of a voxel to one feature is related to its sensitivity to the other feature, ignoring the direction of encoding. Pure selective encoding predicts a negative relationship, mixed selective encoding predicts no relationship, and sparse mixed selective encoding predicts a positive relationship. Similarity matrices averaged over 10,000 simulations. **D)** Correlating the absolute encoding weights, we found that the IPS profile was consistent with sparse mixed selective encoding.

**Figure S7.**
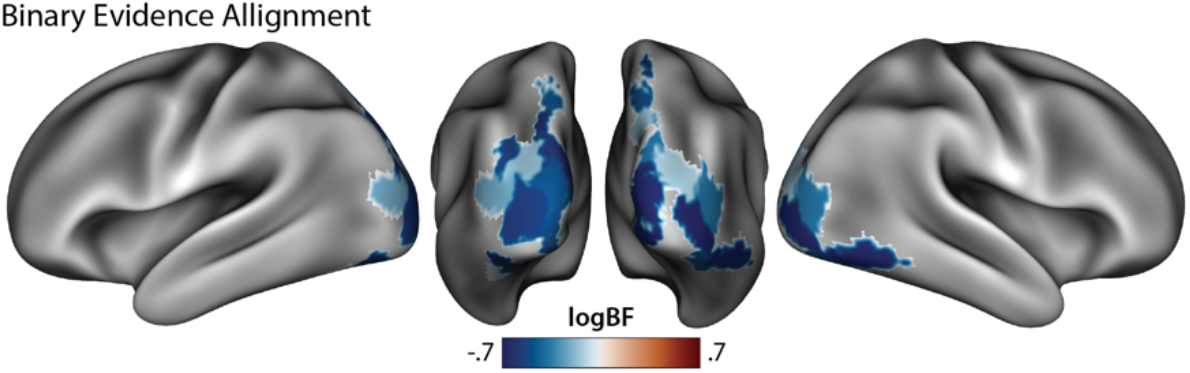
*Binary evidence encoding control analysis.* Target-distractor response encoding alignment using binary evidence rather than coherence-modulated evidence. Note the similarity to Figure 4D.

**Figure S8.**
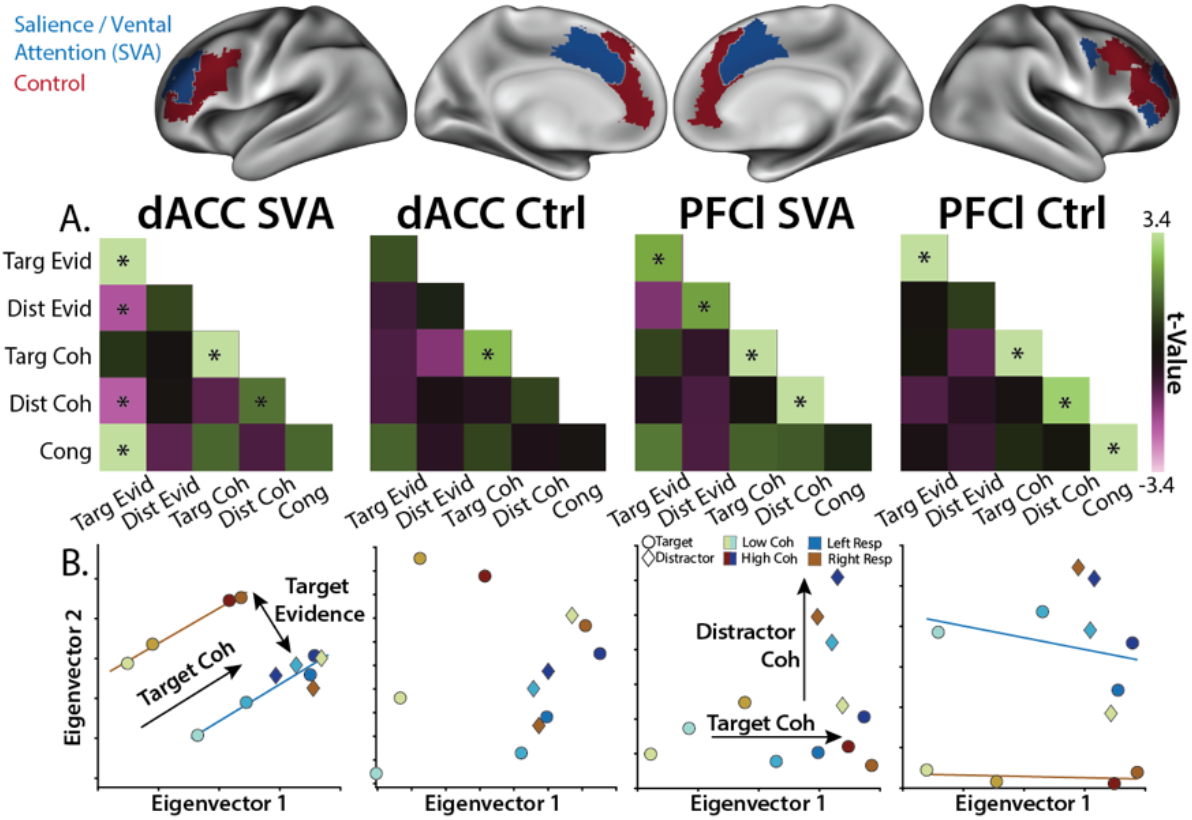
*Feature encoding in frontal networks.* **A)** Similarity matrices for ‘Salience / Ventral Attention (SVA)’ and ‘Control’ networks in dACC and lPFC, correlating feature evidence (‘Evid’), feature coherence (‘Coh’), and feature congruence (‘Cong’). Encoding strength on diagonal (right-tailed *p*-value), encoding alignment on off-diagonal (two-tailed *p*-value). **B)** Classical MDS embedding of target (circle) and distractor (diamond) representations at different levels of evidence. Colors denote responses, hues denote coherence. GLMs: A: Feature MV, B: Evidence Levels, see Table 2.

**Figure S9.**
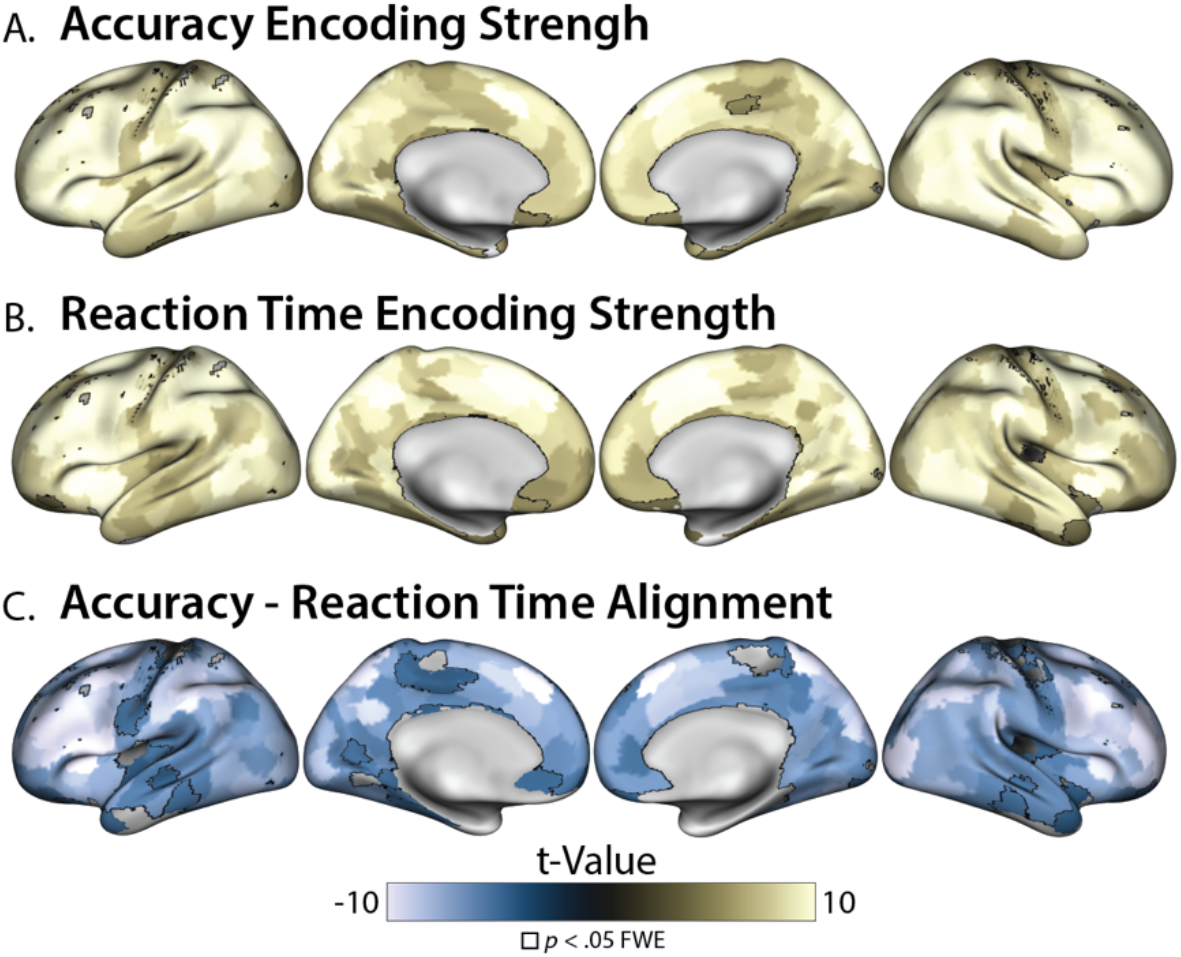
*Performance encoding.* Encoding Strength (across-run reliability) for **A)** Accuracy and **B)** Reaction Time (B). **C)** Alignment between Accuracy and Reaction Time encoding. Outlined parcels are significant at *p* < .05 FWE (max-statistic randomization test). Parcels in C are thresholded based on the reliability in A and B (both *p* < .001). GLM: Performance.

**Figure S10.**
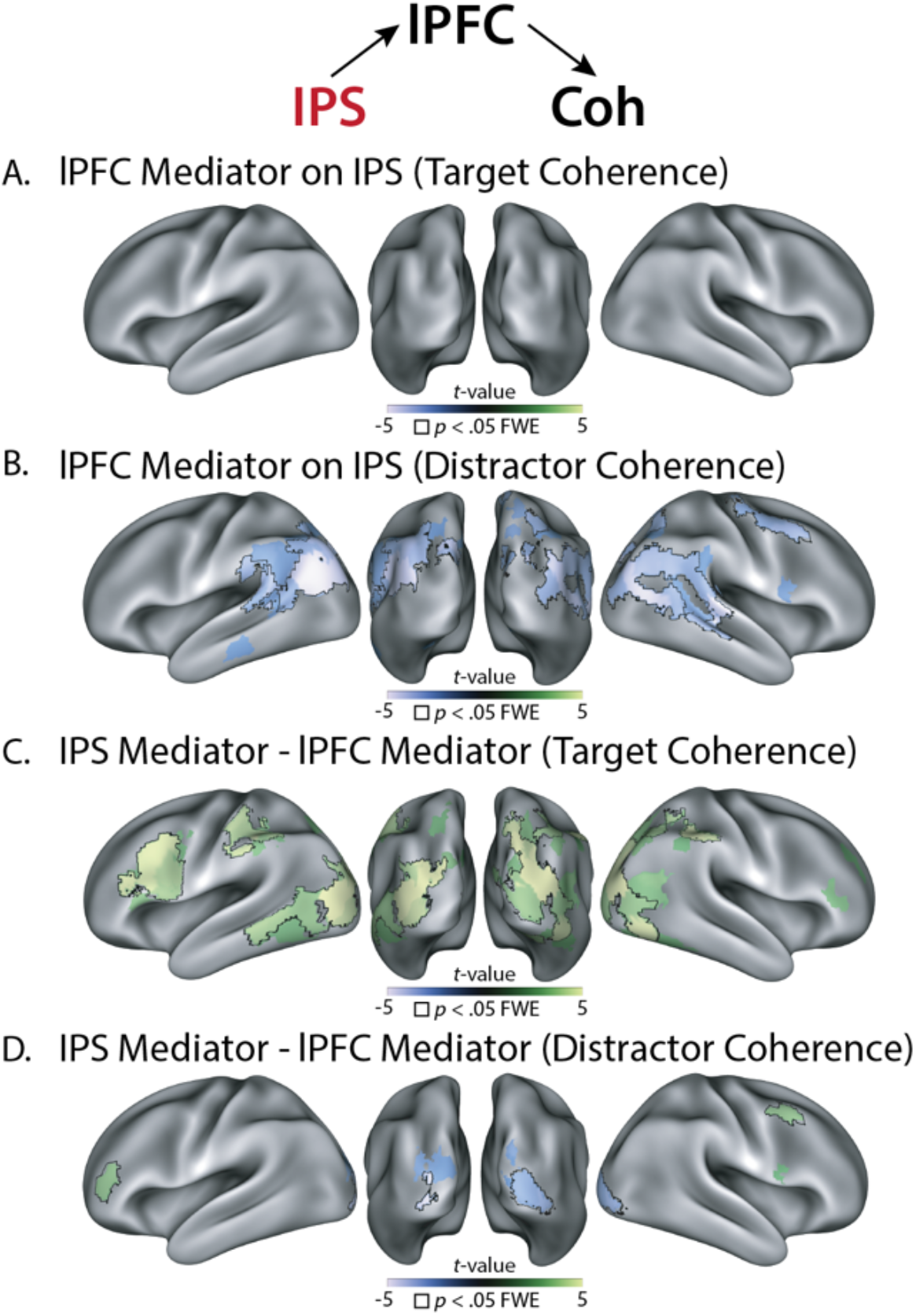
*lPFC mediation.* IPS→lPFC→Coherence mediation for target coherence (A) and distractor coherence (B; compare to Figure 7c). Contrast between IPS-mediation and lPFC-mediation for target coherence (C) and distractor coherence (D).

**Figure S11.**
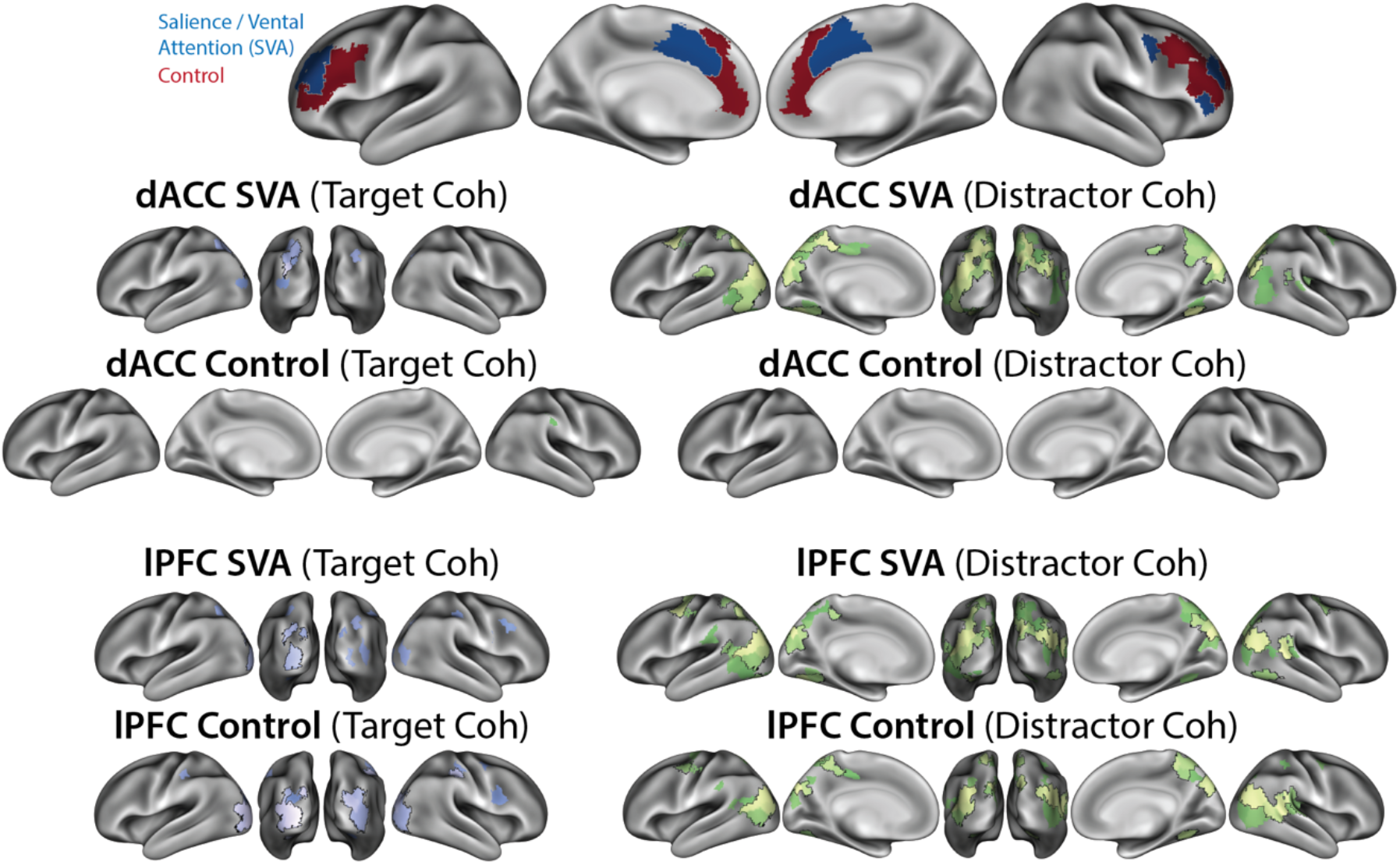
*Coherence alignment with frontal networks.* Activity in ‘Salience / Ventral Attention (SVA)’ and ‘Control’ networks within dACC and lPFC (rows), aligned with target and distractor coherence (columns). Note the similarity between dACC SVA parcels and lPFC parcels.

**Figure S12.**
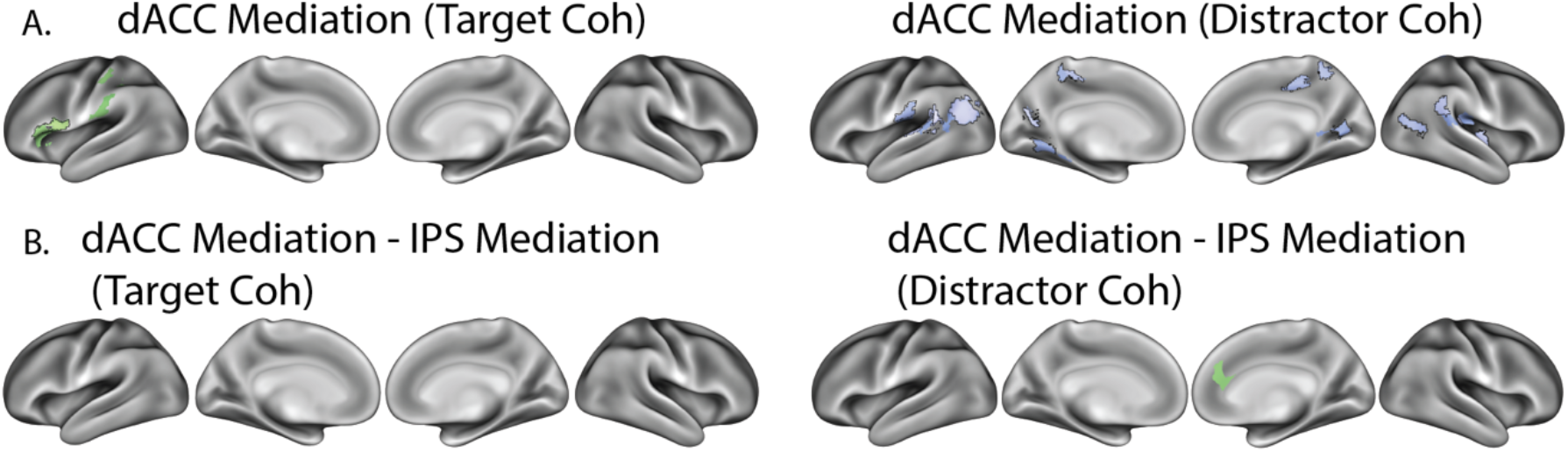
*IPS mediation of dACC connectivity.* **A)** IPS mediation of dACC connectivity (difference in dACC-coherence alignment with and without including IPS predictors). B) Difference between ‘IPS mediation of dACC’ and ‘dACC mediation of IPS’. The lack of activation suggests that this relationship is bidirectional, or originates from a common cause. dACC seed is from the ‘Salience / Ventral Attention’ network (see Supplementary Figure 11).

**Figure S13.**
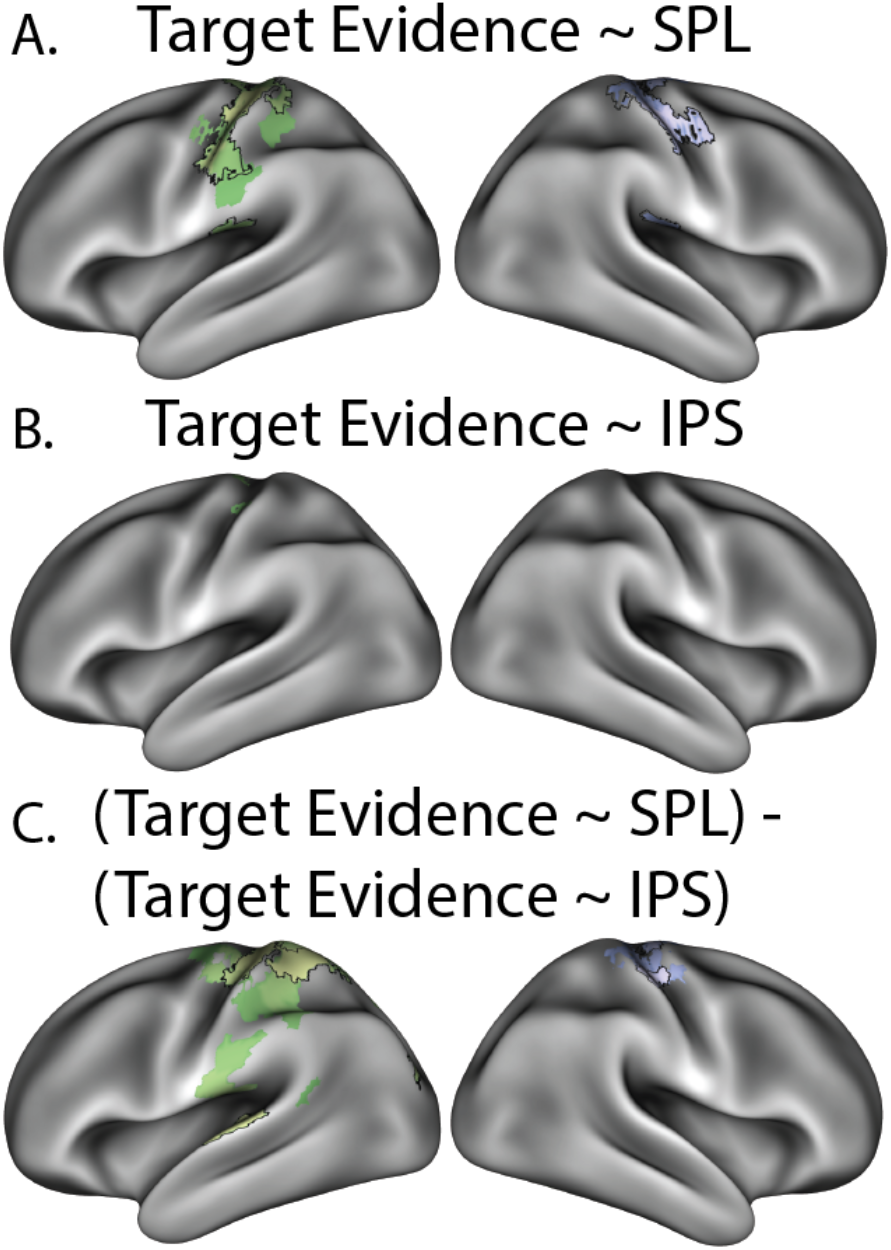
*SPL alignment with evidence encoding.* **A)** Alignment between SPL activity and target evidence encoding. **B)** Alignment between IPS activity and target evidence encoding. **B)** Differences between SPL-evidence alignment and IPS-evidence alignment, showing stronger SPL connectivity. Note that target evidence encoding is signed according to the right-hand response (contralateral motor cortex should have a positive response).

**Figure S14.**
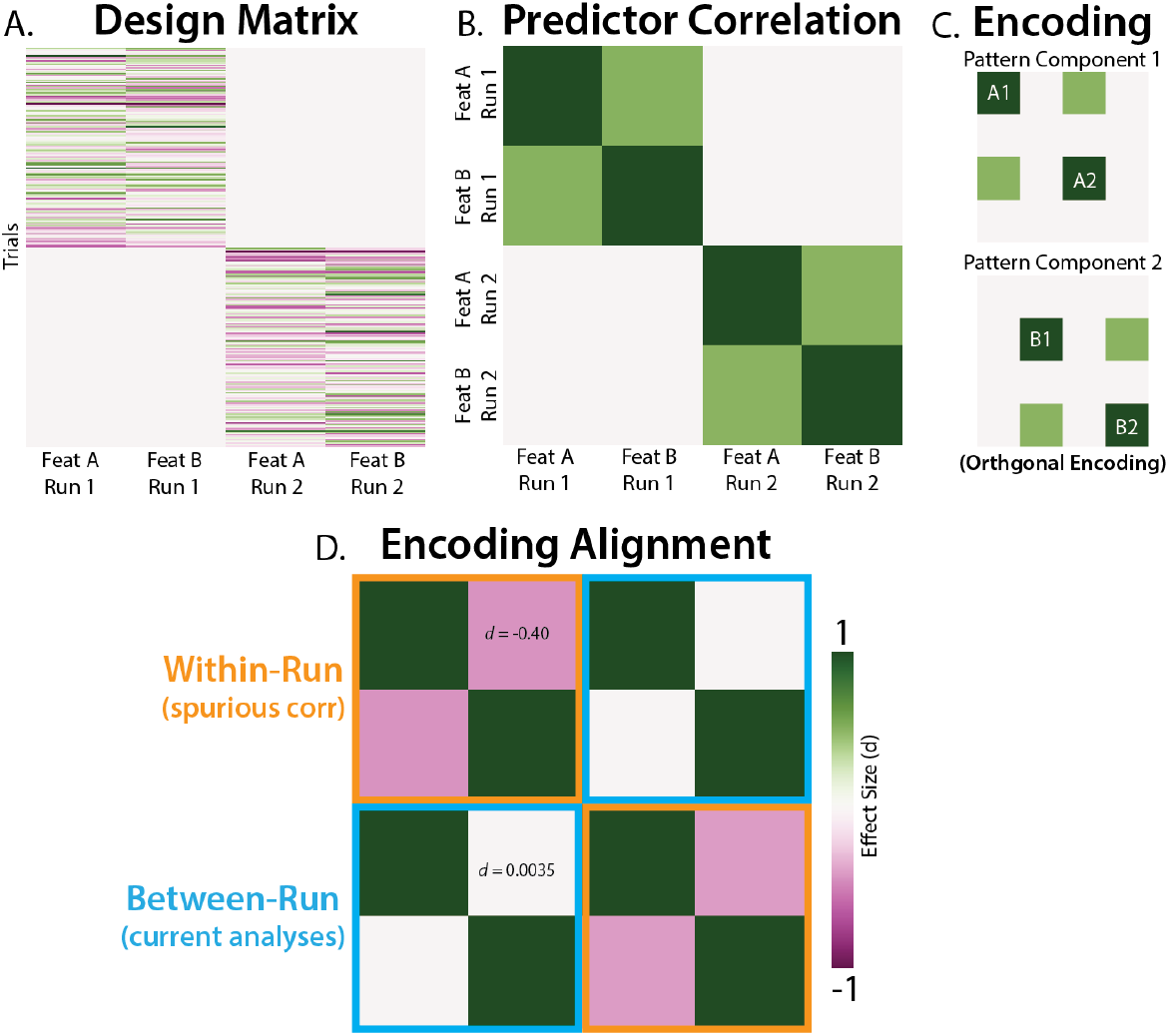
*Cross-validation avoids feature correlations biasing alignment.* We used pattern component modeling (Diedrichsen et al., 2018) to simulate neural data, testing whether feature correlations could spuriously create encoding alignment. **A)** Our design matrix had two simulate runs of two feature timeseries. **B)** Our features were correlated by design (i.e., the columns of the design matrix were correlated). **C)** Despite correlation in the design matrix, these features were independently encoding in our simulated neural population (i.e., in two distinct pattern components, which were each reliable across runs). **D)** Correlating our estimated encoding profiles, we found that within-run alignment (orange) had a spurious negative correlation (the opposite direction of the feature correlations). Critically, our analyses used between-run alignment (cyan), which avoids this biasing effect of feature correlations. Intuitively, since features are not correlated across runs (i.e., they come from different trials), they do not produce spurious correlations. Effect sizes are computed across 10,000 simulations.

**Table S1.** *Partial correlations between coherence and performance*. Correlations between individual differences in coherence-performance alignment, controlling for coherence and performance encoding reliability. Since reliability determines alignment (Spearman, 1987), similarity in alignment may be confounded with similarity in reliability. Overall, these results are qualitatively similar to the zero-order correlation (see Figure 6), albeit with weaker correlations for target coherence. These correlations are particularly robust in IPS.

| Correlation | Covariates | dACC | IPFC | SPL | IPS |
| --- | --- | --- | --- | --- | --- |
| Target-Accuracy,<br>Target-RT | Target,<br>Accuracy, RT | $r_{(27)} = -0.32$<br>$p = .11$ | $r_{(27)} = -0.36$<br>$p = .067$ | $r_{(27)} = -0.11$<br>$p = .56$ | $r_{(27)} = -0.47$<br>$p = .017$ |
| Distractor-Accuracy,<br>Distractor-RT | Distractor,<br>Accuracy, RT | $r_{(27)} = -0.71$<br>$p = 0.50 \times 10^{-4}$ | $r_{(27)} = -0.43$<br>$p = .027$ | $r_{(27)} = -0.48$<br>$p = .012$ | $r_{(27)} = -0.59$<br>$p = .0014$ |

